# Exploring the design space of recombinase logic circuits

**DOI:** 10.1101/711374

**Authors:** Sarah Guiziou, Guillaume Pérution-Kihli, Federico Ulliana, Michel Leclère, Jérôme Bonnet

## Abstract

Logic circuits operating in living cells are generally built by mimicking electronic layouts, and scale-up is accomplished using additional layers of elementary logic gates like NOT and NOR gates. Recombinase-based logic, in which logic is implemented using DNA inversion or excision, allows for highly efficient, compact and single-layer design architectures. However, recombinase logic architectures depart from electronic design principles, and gate design performed empirically is challenging for an increasing number of inputs. Here we used a combinatorial approach to explore the design space of recombinase logic devices. We generated combinations and permutations of recombination sites, genes, and regulatory elements, for a total of ~19 million designs supporting the implementation of all 2- and 3-input logic functions and up to 92% of 4-input logic functions. We estimated the influence of different design constraints on the number of executable functions, and found that the use of DNA inversion and transcriptional terminators were key factors to implement the vast majority of logic functions. We provide a user-friendly interface, called RECOMBINATOR (http://recombinator.lirmm.fr/index.php), that enable users to navigate the design space of recombinase-based logic, find architectures implementing a specific logic function and sort them according to various biological criteria. Finally, we define a set of 16 architectures from which all 256 3-input logic functions can be derived. This work provides a theoretical foundation for the systematic exploration and design of single-layer recombinase logic devices.

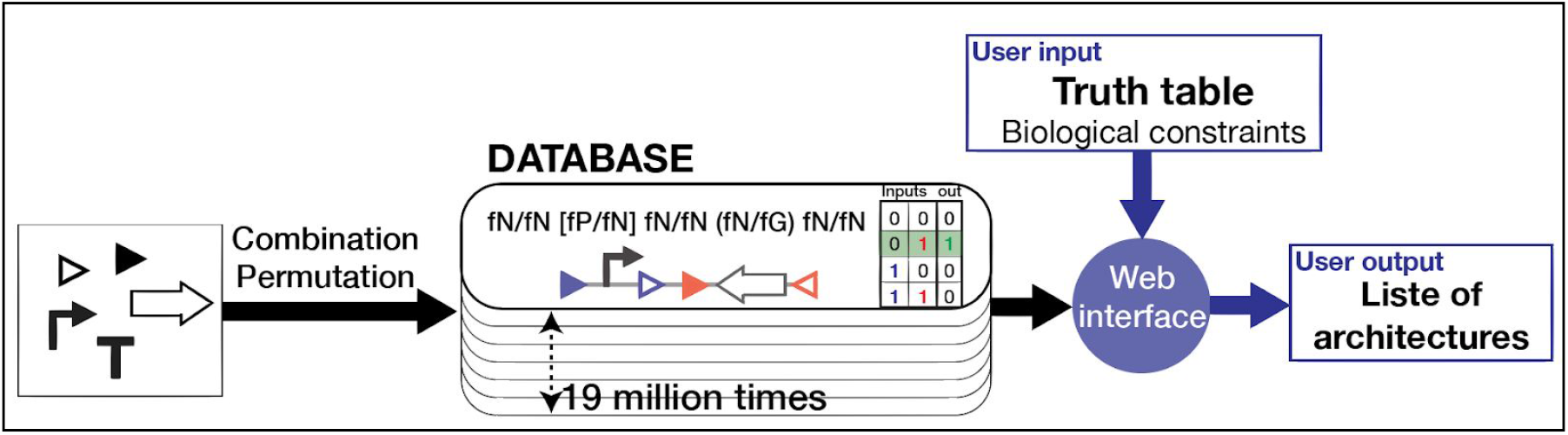

## INTRODUCTION

The field of synthetic biology aims at engineering new biological systems and functions to solve pressing challenges in health, environment, or manufacturing, and to answer basic research questions ^1^. Most engineered biological systems are designed using sensor, signal processing, and output modules ^2 3^. In the past years, large efforts have been devoted to reprogram cellular behavior by engineering logic devices operating within living cells using transcriptional regulators ^4,5^, RNA molecules ^6,7^, proteins ^8,9^, or site-specific recombinases ^10–12^.

Recombinase logic is of particular interest as it supports the implementation of complex logic functions using reduced, single-layer designs ^10^. Recombinase logic devices operate via recombinase mediated irreversible inversion or excision of regulatory elements controlling gene expression. Recombinase logic devices are highly modular (i.e. inputs are easily modified by changing the control signal driving integrase activity), are capable of data storage ^13–15^, and can be adapted to various species with minimal modifications ^11,16–19^.

Several recombinase logic systems operating in single-cell or multicellular systems have been described in recent years, mostly using serine integrases ^10–12^. All 2-input logic functions have been realized through Boolean Integrase Logic (BIL) gates, using a combination of DNA inversion, excision, asymmetric terminators and promoters, and one pair of sites and integrase per input ^10^ (Table S1). Another approach, termed BLADE, used excision-based recombinase devices and integrase site variants to implement all 2- and 3-input logic gates in single-layer single-cell devices ^11^. BLADE offers a single, modular genetic layout albeit with a less compact design since the number of required integrase site variants increases exponentially in the number of inputs. The BIL gate strategy on the other hand is more flexible and can potentially lead to most compact genetic layouts, which could be easier to debug, implement, and scale, while however being highly divergent and *ad-hoc*.

Here we aimed to explore the design space of recombinase logic devices built using the BIL gate strategy. Other types of cellular logic (e.g. transcription-factor based) have explored circuit design spaces using electronic principles ^4^. Recombinase logic devices however mark a departure from electronic layouts mimikry, and methods to explore their design space have been lacking. Systematic design rules have been defined to generate a reduced set of recombinase logic devices ^20^, but these approaches do not support the design of all possible single-layer recombinase logic devices.

In this work, we set up two objectives: (i) to systematize the design of recombinase logic devices based on the BIL gates strategy (Table S1) for an increasing number of inputs (i.e 3 and 4 inputs), and (ii) to determine if all 3-input and 4-input logic functions could be implemented using BIL gate design.

We used a combinatorial approach in which we generated million of combinations and permutations of recombination sites, genes, and regulatory elements. The ~19 million architectures generated in this work can implement all 2- and 3-logic functions and 92% of 4-input logic functions. We provide a web-interface, called RECOMBINATOR, from which users can obtain all possible architectures for any desired logic functions, and sort them according to specific biological constraints. Finally, we defined a reduced set of sixteen logic functions and corresponding architectures which once optimized should support the implementation of all 3-input logic functions. The RECOMBINATOR database supports the exploration of a wide range of single-layer recombinase-logic circuits designs and will facilitate the deployment of recombinase logic for many synthetic biology applications.

## RESULTS

### 1 Formalizing logic architecture generation

Recombinase-based logic devices are built by composing recombinase sites with biological parts controlling gene-expression, plus at least one gene to provide the device output. In order to generate our library of logic designs, we first developed a formalization for biological parts functionality (semantics to which are associated biological elements) (Supplementary text 3.1). We then generated recombinase site arrays (structures) that were functionalized by combinatorially inserting semantics at each intersite position. These functionalized structures were then converted to architectures by replacing semantics by their corresponding biological element (Fig S1).

#### Formalizing functionality of biological parts

In our design, we used three types of biological parts: promoter, terminator, and gene. While these parts for gene expression can be composed in an infinite manner, the parts and the composition of these parts are reducible to a limited number of semantics corresponding to their function in the context of gene expression. We formalized here this finite set of semantics.

Each biological part implement one semantic: a promoter promotes the initiation of transcription (fP), a terminator terminates transcription (fT), and a gene encodes for a function (encoded by a RNA molecule and/or a protein) (fG) (Fig1C). We assume that the semantic of these parts is neutral in reverse orientation (fN). Each part has consequently a forward and reverse semantic (f−/f−), e.g. a promoter in forward orientation has a fP semantic and a fN semantic in reverse orientation, i.e. (fP/fN) (Fig1D).

**Figure 1:**
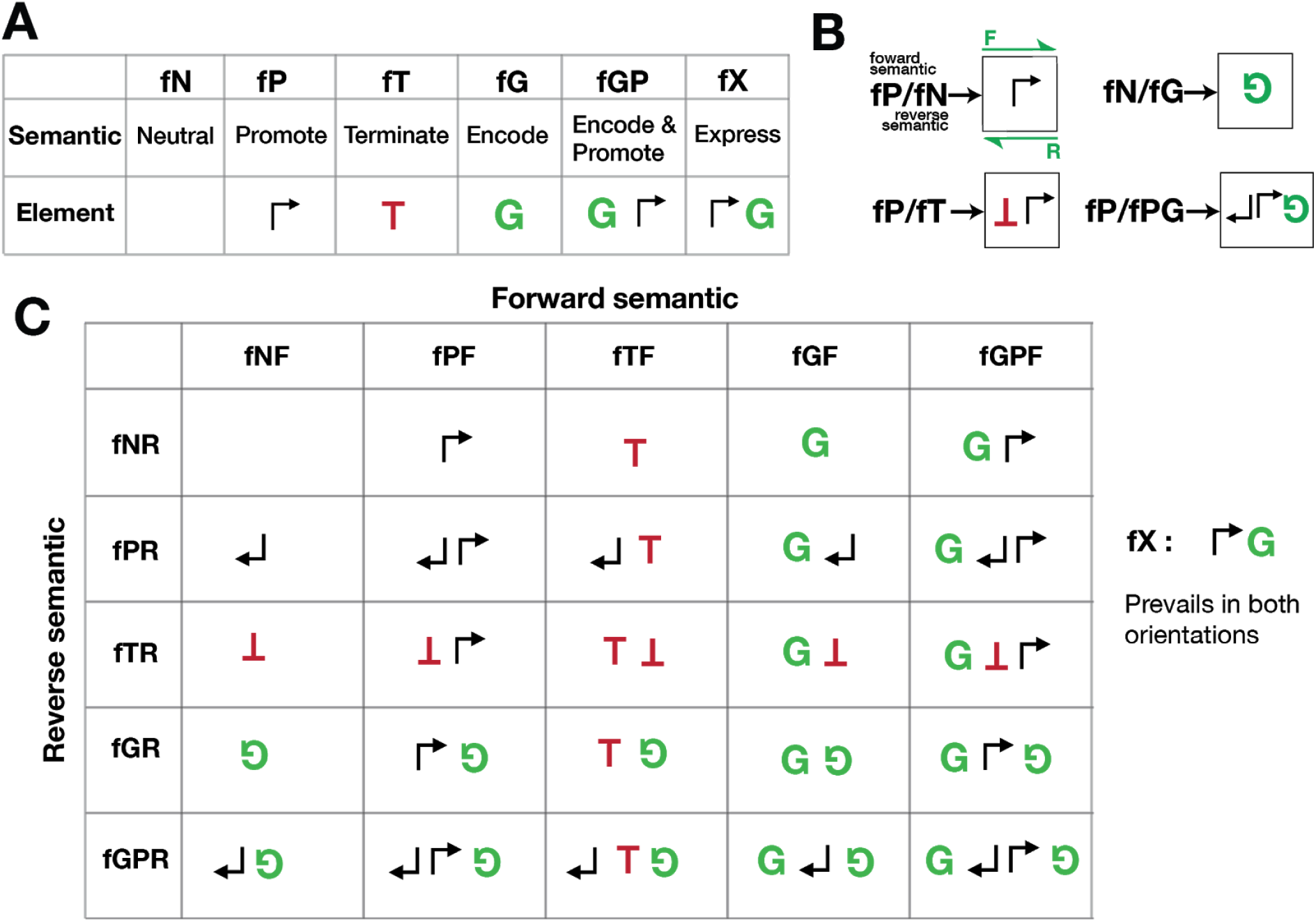
Formalizing functionality of biological parts and part concatenations into a reduced set of semantics. **(A) Formalism for the 6 semantics for a single DNA orientation and corresponding biological elements.** In terms of gene expression, DNA parts and their composition can have 6 different functionalities in a single DNA orientation, called semantics: neutral (no function), promote (promoter), terminate (terminator), encode (gene coding sequence), encode and promote (concatenation of gene followed by a promoter), and express (concatenation of promoter then a gene). **(B) Example of forward and reverse semantics for parts and part concatenations.** Each part and part concatenation have a semantic in forward and in reverse orientation represented as f(forward)/f(reverse). **(C) Correspondence between semantics and elements.** The columns correspond to the forward semantics and the lines to the reverse semantics. In each cell of the table, the element corresponding to semantics concatenation is represented. As the “express” semantic (fX) prevails over any other semantic in both orientations, it is represented separately at the right of the table.

The combination of parts two-by-two leads to only two novel semantics, as several combinations are otherwise being simplifiable in the four previously defined semantics (Table S2). The two novel semantics are: (1) the expression of a gene (fX) corresponding to the concatenation of a promoter with a gene, and (2) the double semantic encapsulating both promotion and encoding (fGP) corresponding to the concatenation of a gene with a promoter (Fig1C).

To summarize, in one orientation (e.g. 5’->3’), all concatenations of biological parts can be reduced to 6 semantics: neutral (fN), promote (fP), terminate (fT), encode (fG), express (fX), encode and promote (fGP). Considering both orientations, 26 semantics exist, as the semantic “express” prevails over all other semantics in both orientations (Fig1; Table S2; Supplementary text 3.1).

#### Representing recombination site arrays as structures composed of well-balanced sequences of brackets and parentheses

Site-specific recombinases drive the logic devices by catalyzing transitions between different recombination intermediates having different output states. Each input signal induces the activation of a single recombinase using either transcriptional, translational or post-translational activation. A recombinase recognizes two specific DNA sequences called pair of recombination sites and mediates excision of the DNA sequence placed between sites in parallel orientation, or inversion for sites placed in antiparallel orientation (Fig2A-B).

**Figure 2:**
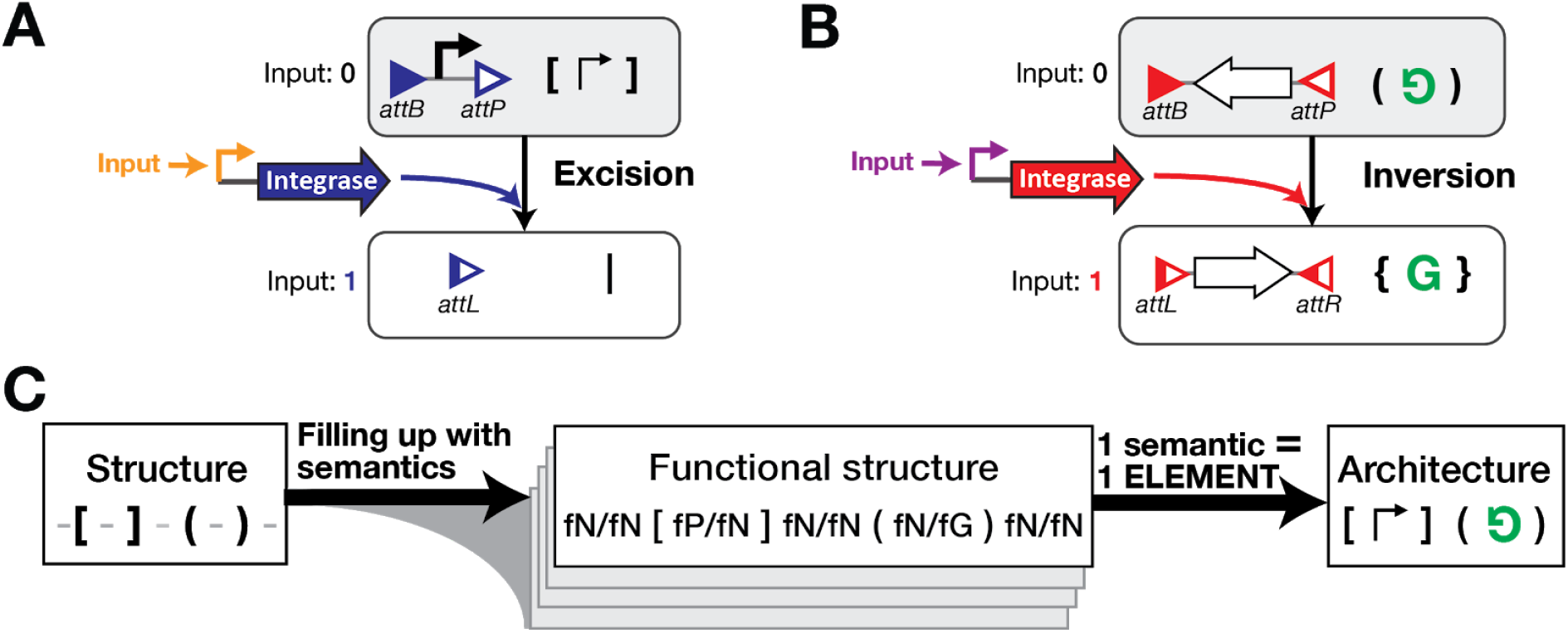
Formalization rules for generating a complete and finite design space. **(A and B) Recombination mechanisms and its formalism.** Each integrase is expressed in the presence of a specific input signal. In A, the integrase sites (triangles) are in the same orientation, the integrase mediates excision of the DNA between the two sites (here a promoter). Brackets are used to represent integrase sites in excision orientation and a vertical bar for used site (either attL or attR) resulting from the excision. In B, the integrase sites are in opposite orientation, inversion of the DNA between the two sites is then performed by the integrase (here a gene). Parentheses are used to represent integrase sites in inversion orientation and curly braces for the resulting used sites (both attL and attR). **(C) Workflow for generating an architecture**. Site structures are generated as functionalized Dyck words. Each space between sites, considered as variable, is filled with semantics using a backtracking algorithm to obtain functionalized structures. Each semantic is associated to one element, so that the architecture corresponding to each functionalized structure is obtained by direct replacement.

In order to generate all possible site permutations, we used brackets and parentheses to represent recombination sites respectively in inversion and excision orientations. We called *structure* the sequence of parentheses and brackets representing a recombination site array. As we focused on combinatorial logic, recombination reactions must be independent, therefore recombination sites must not be interleaved. We thus generated only structures composed of well-balanced sequences of brackets and parentheses (corresponding to a Dyck word, see methods and supplementary materials for details). Consequently, the identification of site pairs is unambiguous, and no annotation of parentheses and brackets is required (e.g. Fig S2).

#### Architecture generation

To implement logic, biological parts are placed between integrase sites conditioning their excision or inversion and subsequent effect on gene expression. Therefore, after having generated structures, we generated *functional structures* by placing a semantic in each available space between brackets or parentheses.

While various biological part concatenations can have the same semantic, we selected a single element to encode each of the 26 semantics (Fig1C-D). The choice of each element was guided by several criteria. As a general design rule, the chosen element had to be the simplest concatenation of parts capable of encoding the semantic of interest. More specifically, we aimed to use elements composed of the minimum number of parts, and respecting as much as possible the following two architectural constraints (e.g. Fig S3A). First, we prioritized avoiding promoters facing each other, a configuration known to generate interferences and unexpected transcriptional behavior (e.g. PR-PF is favored to PF-PR) ^21,22^. Then, we chose elements having the minimum number of parts between a gene and the promoter controlling its transcription (e.g. GF-TR is favored to TR-GF), as gene expression generally decreases with the distance between the promoter and the gene ^23^.

We then generated architectures from functional structures by replacing semantics by their corresponding element (Fig2C).

### 2 Algorithmic method for obtaining irreducible and non-redundant architectures

We aimed at generating a recombinase device database composed of irreducible and non-redundant architectures and without chiral pairs of architectures. An irreducible architecture is an architecture in which no part can be removed without changing the logic implemented (e.g. Fig S3B). A non-redundant architecture is an architecture which implement a Boolean function which is not simplifiable into a Boolean function with a reduced number of inputs. A chiral pair of architectures corresponds to two architectures which are reverse-complement from each other (e.g. of chiral structures Fig S2C). We performed the generation and sorting at the level of functionalized structures, and the obtained set of irreducible and non-redundant functionalized structures was then converted in architectures (Fig S1).

From one architecture, we generated the derived architectures, corresponding to the different architectures resulting from integrase-mediated recombination (Fig 3A). A derived architecture corresponds to a specific input state, and from one architecture: 2^N-1 derived architectures are generated, with N being the number of integrase site pairs (equivalent to number of inputs). For each input state, the algorithm determines the gene expression status of the architecture. Determination of gene expression status was performed using a set of rules (Supplementary text3.3 and Table S2) that translate the total concatenation of semantics into a general semantic indicating its gene expression state, enabling the generation of the architecture’s truth table.

**Figure 3:**
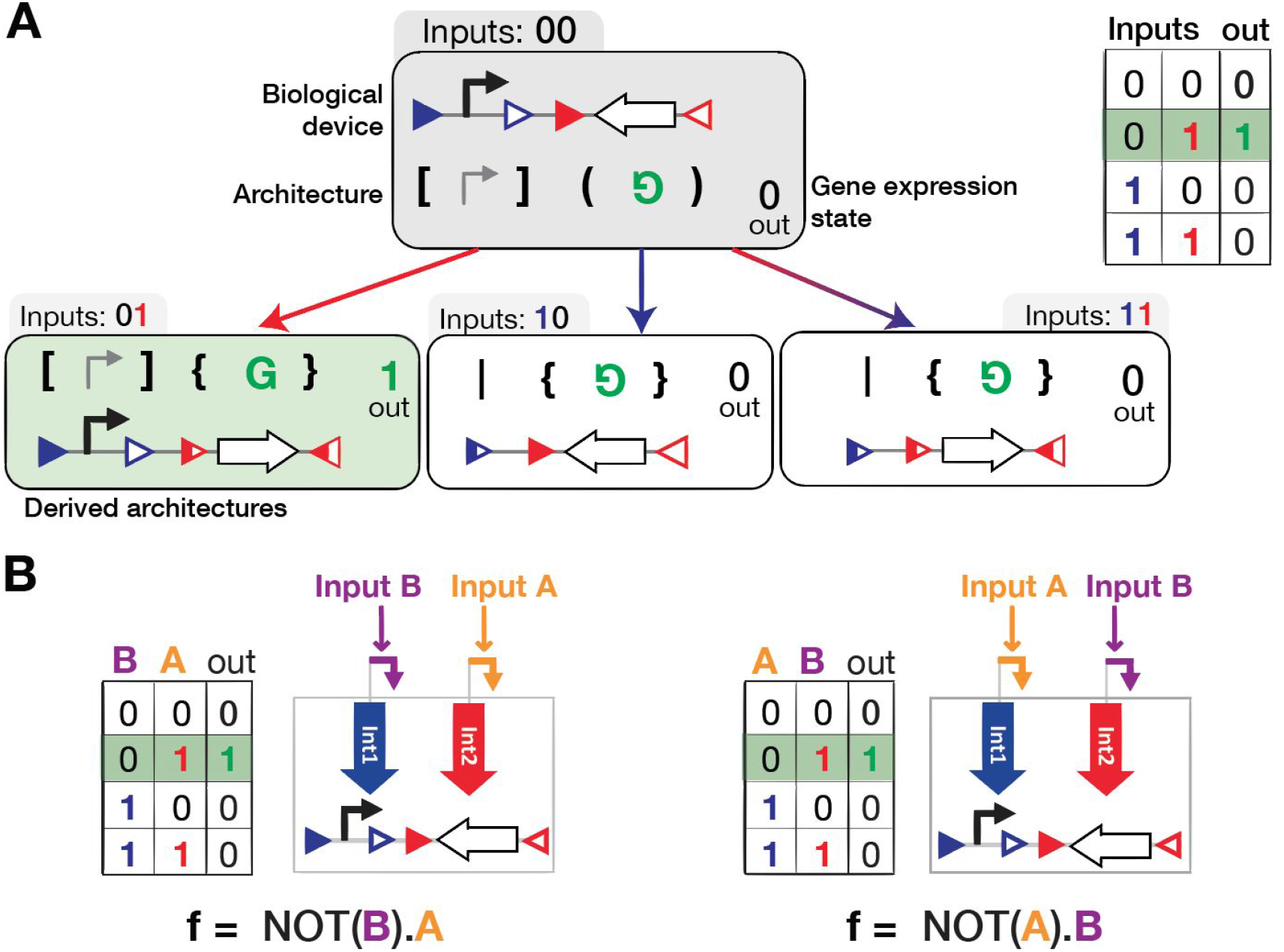
From architecture to truth table. **(A) Generation of architecture derived states and of the corresponding truth table.** For a given architecture, here also represented as a biological device, the derived architectures corresponding to the different input states are obtained by simulating integrase recombination. In this example, in the presence of one input (red) corresponding to the state 01, the gene is inverted, in the presence of the blue input (state 10), the promoter is excised and in the presence of the two inputs, the promoter is excised and gene is inverted. From each architecture and derived architecture, the gene expression state is obtained, and the truth table is generated. **(B) From one architecture, implementation of several logic functions belonging to the same P-class.** By differentially connecting inputs to integrases, various logic functions are obtained from the same architecture. For example, the function NOT(B).A realized by connecting input B to integrase 1 (blue) and input A to integrase 2 (red). The permutation function NOT(A).B is obtained by permuting the connections.

Importantly, during the generation, we did not attributed inputs to integrases and recombination site pairs. All parenthesis and/or brackets can eventually be associated to a specific input. All logic functions corresponding to the various input permutations are thus implementable with one architecture (Fig3B). As logic functions are in the same permutation-class (P-class) if they are equivalent by permutation of inputs, a given architecture implements a complete P-class ^24^.

### 3 Database analysis

We performed the database generation for up to 4 inputs. We obtained 18,163,227 architectures implementing 3,608 P-classes corresponding to 59,820 logic functions. We found that using our specifications, all 1-, 2- and 3-input functions and P-classes were implementable (Table 1). On the other hand, 92.25% of the 4-input logic functions were implementable, corresponding to 90.4% of the 4-input P-classes (the discrepancy between the number of functions and P-classes arise from the fact that P-classes contain a different number of functions).

**Table 1:** Database characteristics. The number of generated architectures and the corresponding functions/P-classes implemented are represented. Only non-redundant Boolean functions are represented. The percentages of implemented functions and of implemented P-classes were calculated as percentages of the total number of functions and P-classes corresponding to a given number of inputs (Table S3).

| Number of inputs | Number of architectures | Number of function implemented | % of implemented functions | Number of P-class implemented | % of implemented P-classes |
| --- | --- | --- | --- | --- | --- |
| 1 | 10 | 2 | 100% | 2 | 100% |
| 2 | 724 | 10 | 100% | 8 | 100% |
| 3 | 96,981 | 218 | 100% | 68 | 100% |
| 4 | 18,065,512 | 59,590 | 92.25% | 3530 | 90.4% |

#### Implementability of logic functions

Over all the 3,904 4-input P-classes, 374 P-classes are not implementable with our circuit specifications. These P-classes are then the more complex ones to implement using recombinase logic. We sought to understand if non-implementable P-classes shared common properties. We started by searching if a correlation existed between the properties of the logic equations and P-class implementability.

We write logic functions as a sum of product of NOT or IDENTITY functions, corresponding to the minimal disjunctive form also called minimal SoP (sum of product) form ^25^. A disjunctive form is called minimal if there exists no other equivalent expression involving fewer products, and no other equivalent expression involving the same number of products but a smaller number of literals ^25^.

We hypothesized that the most difficult P-classes to implement were the ones with the largest equation, such as while written in a minimal disjunctive form, these equations contained the highest number of terms, and the highest number of literals per term. Indeed, the non-implementable 4-input P-classes correspond to the P-class with the highest number of terms and highest number of literal per terms (Fig4A). While a total of 9.6% of 4-input P-classes are not implementable, this number reaches 23.7% for P-classes with more than 4 terms and more than 2.5 literals per term (Table S4).

**Figure 4:**
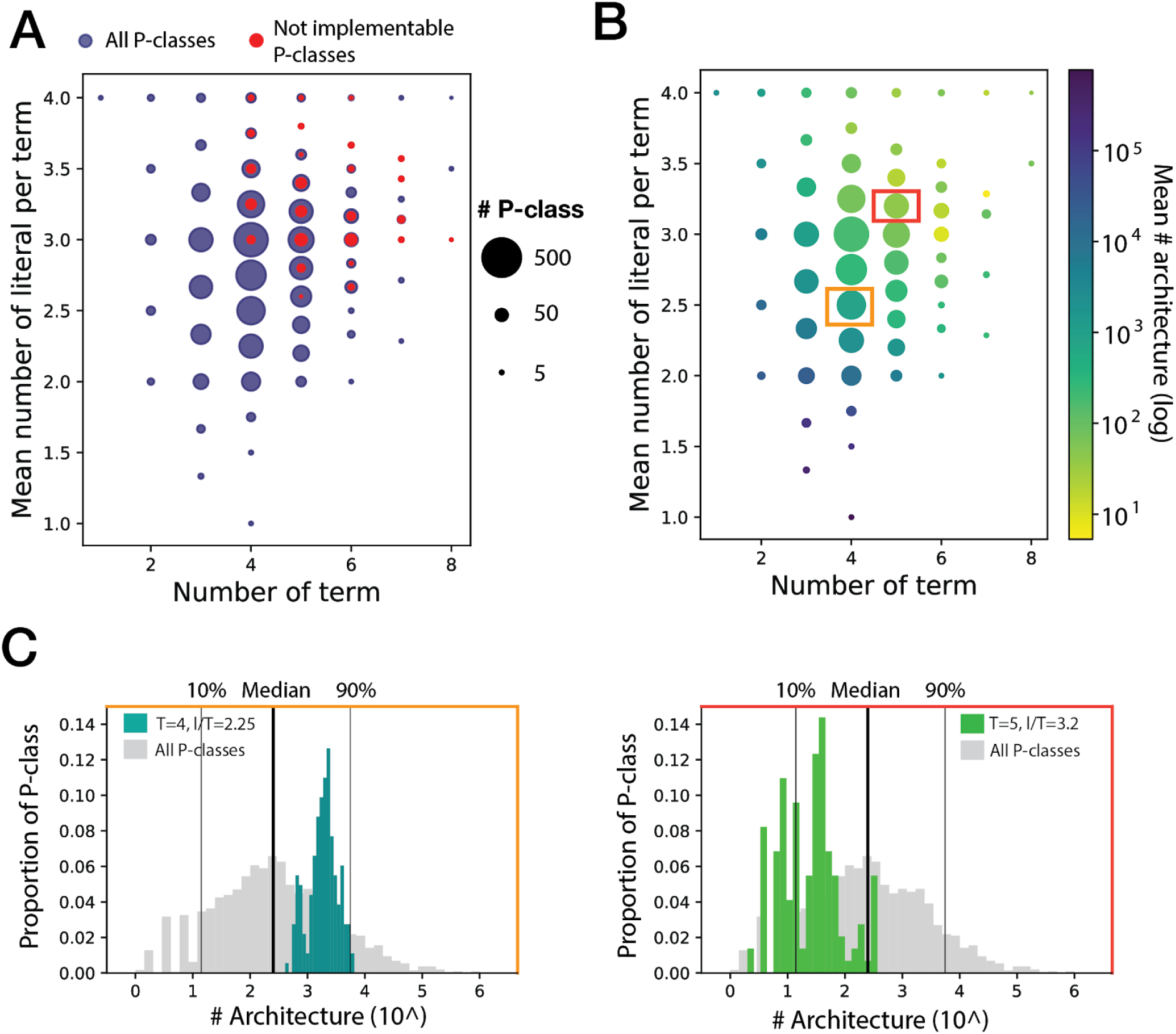
Implementability and number of possible architectures of P-classes according to their complexity. (**A) and (B)** P-classes with a specific number of terms and of literals per term are represented in the corresponding x and y positions by a dot with a diameter proportional in logarithmic scale to the number of P-classes. **Representing in (A) the implementability of P-classes,** the blue dots correspond to all 4-input P-classes and the red ones to the not implementable 4-input P-classes. In (B), only the 4-input implementable P-classes are represented. **Representing the number of possible architectures according the P-class complexity,** the dot colors are associated to the mean number of possible architectures for each set of P-classes. Blue corresponding to a reduced number of architectures and yellow the highest number of architectures. 2 dots are surrounded by respectively an orange and red rectangle, corresponding to the two P-class sets represented in (C). Detailing (B), the distribution of the number of architectures for these two sets of P-classes is represented. For P-classes with 4 terms and 2.25 literals per term, the distribution is in teal (blue/green) and for 5 terms and 3.2 literals per term in light green (used bin is 20). The distribution for all 4-input P-classes is also represented in grey as reference (used bin is 40).

However, we found some exceptions to this trend: some clusters of P-classes with a very high-number of terms and literals per term, such as 8 terms and more than 3 literals per term, and 7 terms and 4 literals per term, were totally implementable. These exceptions correspond to complete or partially symmetric functions (See supplementary text 3.4), such as functions which are entirely or partially dependent on the number of inputs present, e.g. a 2-input XOR function.This can be explained by the fact that, in comparison to other design strategy such as repressor-based logic, symmetric functions are easily implemented with recombinase logic using nested inversion (^10^; Supplementary text 3.4).

#### Number of architectures per P-class

The number of architectures capable of implementing a particular P-class is highly variable, ranging from 1 to 1.4.10^6 (Fig4B) (6 P-classes with only one architecture: Supplementary text 3.5), and widely distributed (Fig4C). For implementable 4-input logic functions, the median number of architectures is of 252, providing a wide design space for most functions.

Similarly than for implementability, P-classes with the highest number of possible architectures are P-classes with the lowest number of terms and literals per term, corresponding to the simplest functions (Fig4B, 4C left panel). Reciprocally, the P-classes with the lowest number of possible architectures are the ones with the highest number of terms and literals per term, corresponding to the more complex functions (Fig4B, 4C, right panel). While the figure 4B represents the mean number of architectures for a cluster of P-classes with a specific number of terms and literals per term, we also plotted for each cluster the distribution of the number of architectures (Fig4C and FigS4). We found that the distribution of the number of architectures was quite homogeneous in each cluster and shifted from a high to a low number of architectures when P-classes complexity increased.

#### Influence of design specifications on P-classes implementability

Previously, various strategies have been defined for the design of recombinase-based devices using either only excision ^11,12,26^ or mainly inversion and terminator based elements ^10^. In our database, we generated all possible designs using both excision and inversion, and a flexible use of the previously-defined biological parts. By sorting these architectures, we obtained the percentage of P-classes implementable with restrictive design criteria corresponding to the previous works implementing recombinase logic ^10,11^.

##### Excision vs inversion

We first analyzed the influence of the type of recombination reaction on P-classes implementability. We found that using only DNA excision highly reduces the number of implementable P-classes (82% of 3-input P-classes implementable and only 26% of 4-input P-classes) (Fig5). In contrast, almost all implementable P-classes can be realized using only inversion (100% of 3-input and 86% of 4-input P-classes) (Fig5). One explanation is that excision is a destructive mechanism leading from a specific semantic to a neutral semantic, reducing the semantic space of the device. On the other hand, inversion leads to a semantic change therefore permitting to implement more complex P-classes (Fig S5A-B). Very interestingly, while all 3-input P-classes are implementable with only inversion, we found that some 4-input P-classes (4.5% of implementable P-classes) require both inversion and excision to be implementable.These P-classes cluster in the medium complexity region of the plot (Fig S5C). Therefore, inversion only is not sufficient to implement the maximum range of Boolean functions.

**Figure 5:**
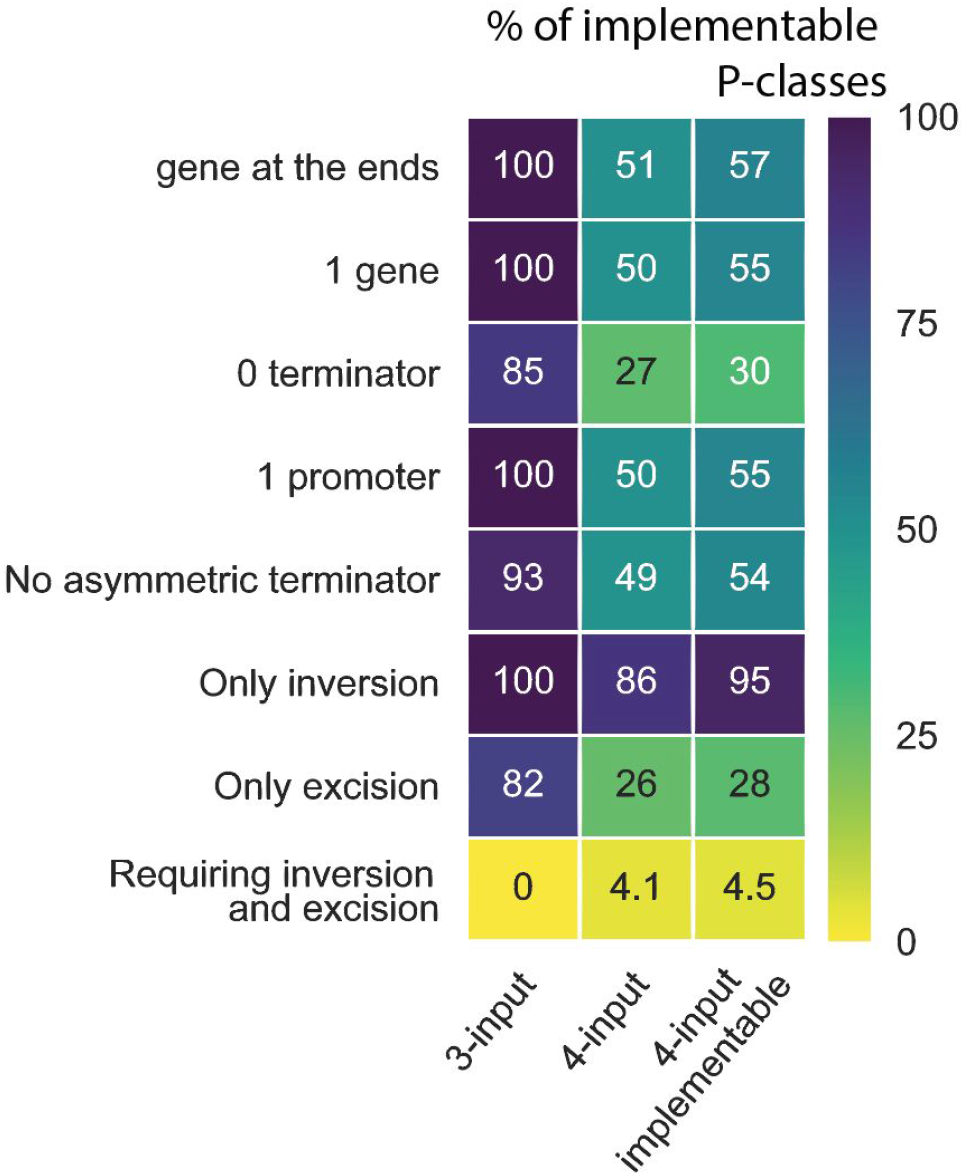
Percentage of P-classes implementable as a function of various restricting criteria. The columns denote the different set of P-classes, such as 3-input, 4-input and 4-input implementable P-classes. Each line corresponds to the percentage of P-classes implementable with the corresponding design specification, the color scale represent their respective percentages, from yellow (0%) to dark blue (100%).

##### Number of genes, promoters, and terminators

A functional logic device requires at least one promoter and one gene, and can therefore be designed without a terminator. However, without terminator, 15% of 3-input P-classes are not implementable, as well as 30% of 4-input implementable P-classes (Fig5). Terminators are thus essential elements for recombinase logic implementation, in particular for the more complex P-classes (Fig S6A). Similarly, by limiting the number of genes or the number of promoters to one, the number of implementable P-classes is reduced by almost 50% for 4 inputs (Fig5). To conclude, using all gene-expression parts and letting flexible their respective numbers permit to maximize the number of implemented logic functions.

In our design, we considered terminators as asymmetric terminators (terminating transcription in a single orientation). Bidirectional terminators (semantic fT/fT) were obtained by combining two terminators in opposite orientations. A reduced number of asymmetric terminators has been characterized, and only in bacteria ^27^; such parts might thus limit the portability of our devices to other organisms, particularly in eukaryotes. We wondered how many P-classes required strictly asymmetric terminators to be implementable. Excluding asymmetric terminators from the designs, we found that 93% of the 3-input P-classes and 54% of the 4-input implementable P-classes were implementable (Fig5 and FigS6B).

We previously designed 2-input logic gates in which a single output gene is placed in 3’ of the device, and is easily interchangeable ^10^. In addition, this design allows the logic device to be directly placed upstream of endogenous genes to add an externally-controlled layer of logic to their regulation. By filtering the database for architectures possessing a single gene in the 5’ or 3’ extremities, we found that all 3-input P-classes and 57% of 4-input implementable P-classes are implementable using this design.

Additionally, we tested the effect of combining criteria two by two, and found that for stringent criteria such as the use of only excision, the combination with another criteria drastically reduced the number of implementable P-classes (Fig S7-8).

Based on literature, we believe that the following criteria should be satisfied in order to obtain well-behaving logic devices:

1. Not having two promoters facing each other, such as architectures respecting the no cross promotion constraint (see material and methods for criteria list).
2. To have a reduced distance between the gene and the promoter leading to the expression of this gene. In addition, in order to minimize differences in output level between the different input states, we recommend:
3. To avoid having two genes expressed at the same time.
4. To avoid having two promoters mediating the expression of one gene.

These criteria are not absolute, and sometimes cannot be all satisfied for implementing all logic functions. Experimental implementation and characterization of various architectures will ultimately be the only way to validate their functionality.

### 4 A web-interface to sort architectures according to user-defined constraints

Based on this large database, we created a user-friendly web-interface allowing the users to obtain all architectures implementing logic functions of their choice (Fig 6). For each architecture, the various specifications of the design are listed, such as the number of inputs, inversions, excisions, genes, terminators, and promoters, and the total number of parts. Additionally, the approximate length of the architecture is specified (calculated by considering as lengths: 20bp for a promoter, 40bp for a terminator and an integrase site, and 1 Kbp for a gene). We also specify if the architecture respects or not design criteria of interest, such as having the gene at the extremity of the construct (5’ or 3’), the absence of asymmetric terminators, or the absence of promoters facing each other. Indeed, while we excluded facing promoters from element design, insertion of elements within functional structures can still result in architectures containing facing promoters. In all cases, the list of architectures can be re-ordered according to a criterion of choice. A filtering tool allows the user to selectively display specific architectures respecting a given set of criteria.

**Figure 6:**
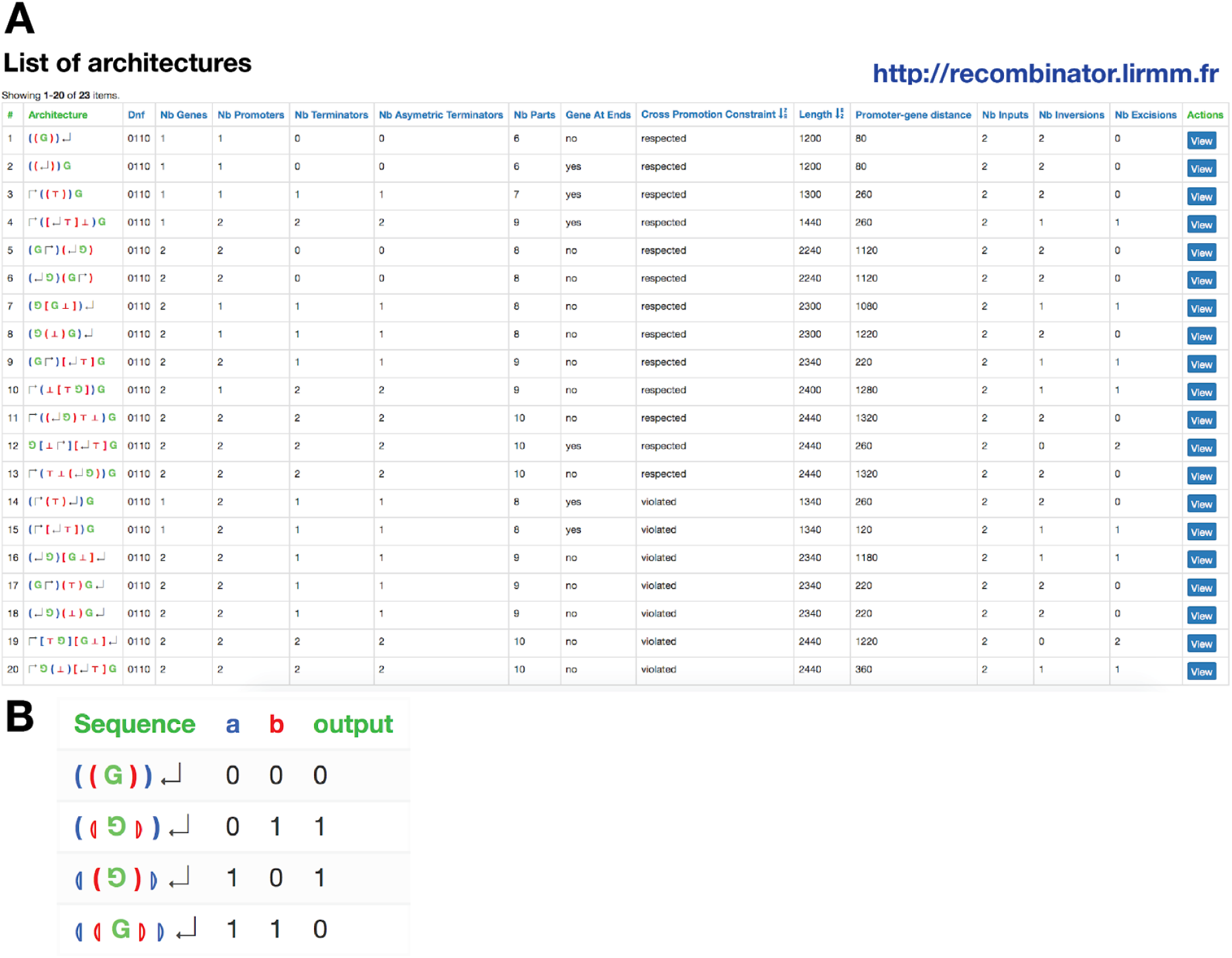
The recombinator web-interface. **A. Example of search results on the recombinator web-interface** with as entry the XOR logic function, 23 architectures are found with here the 20 first architectures. This list can be sorted according to various criteria. **B.** The web-interface also allows the user to directly provide an architecture to obtain all its derived states and the implemented Boolean function.

When the user selects a specific architecture, all the derived states of the architecture are also represented with the correspondence between sites and inputs. Additionally, all logic functions implementable with the same architecture (therefore P-equivalent functions) are accessible from this page.

### 5 A reduced set of 16 NP-equivalent architectures based on DNA inversion to implement all 3-input functions

For most logic functions, the database provides many different possible architectures (with a median of 252, and a maximum of 1.4 million for the logic function in supplementary text 3.6). An important gap remains between theoretical designs and well-behaving biological devices, and experimentally testing all proposed architectures is challenging. We therefore wanted to identify a method enabling us to reduce the number of biological devices to be tested and optimized, while maximizing the number of implementable logic functions. We already used permutation classes to reduce the number of generated architectures by not assignating integrase sites to inputs. Here we built a minimization strategy based of NP-classes, which corresponds to class of logic functions which are equivalent by permutation and negation of inputs ^24^.

Permutation of inputs in integrase-based system is simply performed by permutation of the connection between integrase and inducible promoters. Importantly, negation is very easily performed in recombinase logic by using DNA inversion. For example, the NOT function being the negation of the identity function, a NOT device in recombinase logic can be transformed into an IDENTITY by inverting the asymmetric terminator controlling the flow of RNA polymerase, and vice versa (Fig7A). In this simple case, the negation of the input in a logic function is performed by inversion of the part between integrase sites in inversion orientation. This approach is generalizable to more complex devices that use only DNA inversion. Consequently, starting from any all-inversion architecture, replacing LR sites by BP sites in the recombination intermediates leads to a new architecture implementing a specific P-class within the same NP-class (Fig7B).

**Figure 7:**
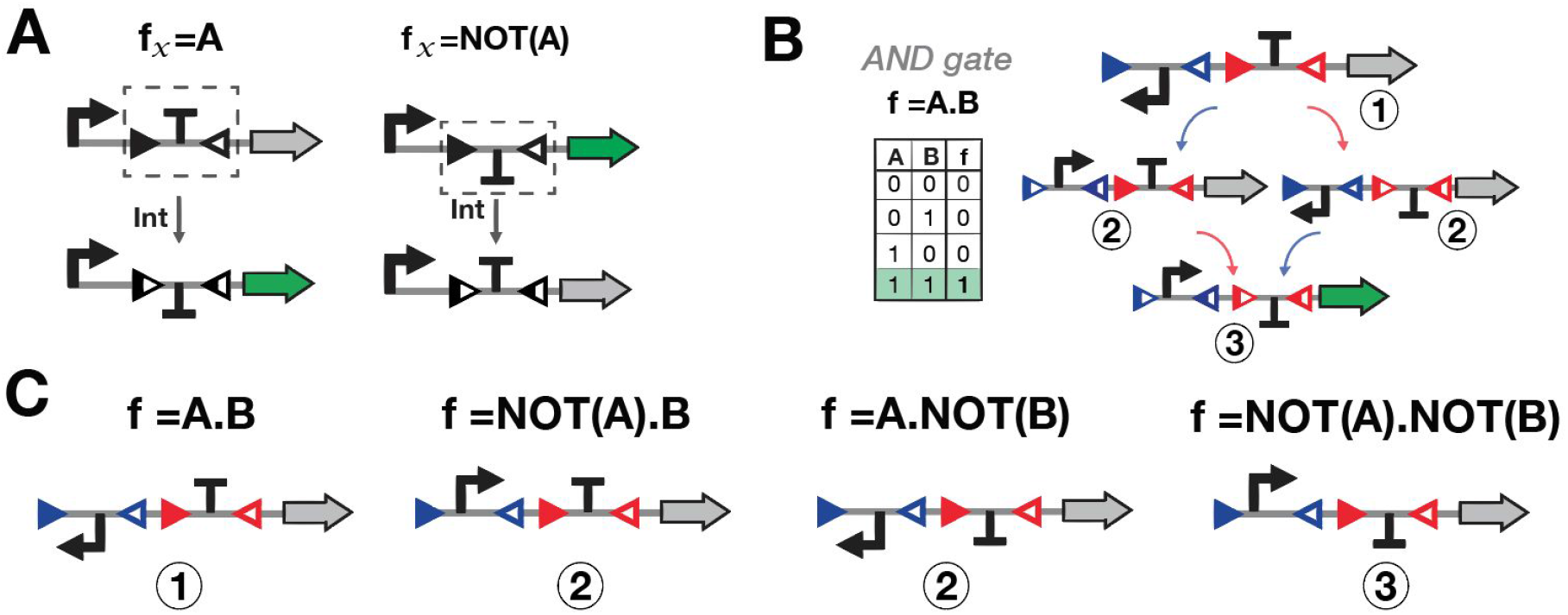
Implementation of NP-equivalent Boolean functions using inversion-based logic devices. (A) IDENTITY- and NOT- logic devices based on the inversion of an asymmetric terminator. (B) A 2-input AND gate based on the combination of promoter and terminator modules and its intermediate recombination states. (C) Implementation of all functions from the 2-input AND gate NP-class. Designs are based on the AND gate represented in B and correspond to the intermediate recombination states in which LR sites have been replaced by BP sites. The inversion of a single part of the device for A.B leads to either NOT(A).B or A.NOT(B), and the inversion of both parts to NOT(A).NOT(B) logic function.

We can thus reasonably assume that for a given NP-class, characterizing in all input states and optimizing a single logic device executing one function should provide a good estimate of the behavior of all derived logic devices implementing the NP-class. 16 and 380 NP-classes exists for 3-input and 4-input, respectively.

Because this approach is restricted to architectures using DNA-inversion, it is theoretically possible to optimize one architecture for each of the 16 three-input NP-classes and for 322 of the 380 four-input NP-classes.

Inversion-only architectures represent only 25.4% of the original set (4.6 million of architectures), thereby leading to a lower probability to find architectures respecting all the biological criteria listed above (Fig S9). We present in Figure 8 an example of architecture for each of the 16 3-input NP-classes, chosen according our recommended criteria.

**Figure 8:**
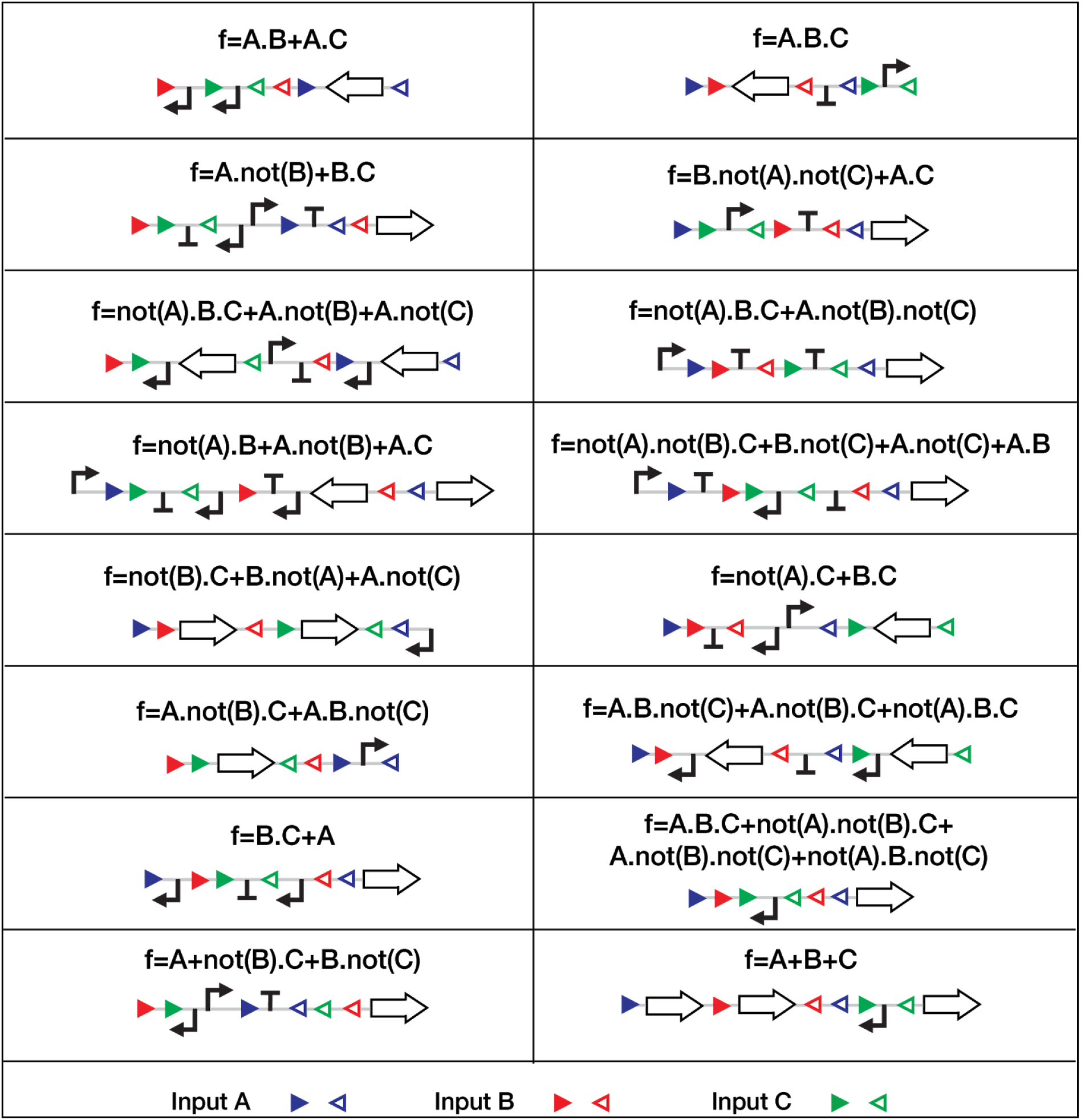
Selected architectures for the implementation of the 16 3-input NP-classes. Each box implements one NP-class. Here one Boolean function and a selected logic device implementing this Boolean function are depicted for each NP-class. Triangles correspond to integrase sites, blue is for variable A, red for B and green for C.

## DISCUSSION

Here we took a systematic approach to explore the design space of recombinase logic devices. We devised a formal language to express and interpret the output state of genetic devices, along with several strategies allowing us to minimize the number of generated constructs to a non-redundant and irreducible set. We obtained a total of ~19 million architectures, to date the largest database of biological logic designs, implementing a total of 59,820 Boolean functions (corresponding to 3,610 P-classes).

We proved that all 3-input and 92% of 4-input Boolean functions are implementable in single cell using integrase-based devices within the constraints of using one integrase and one pair of recombination sites pair per input. Most P-classes can theoretically be implemented by hundreds of architectures. Not surprisingly, the number of possible architectures for a given P-class is inversely correlated with their complexity (i.e. the number of terms and literals per term).

We tested the impact of various design strategies on function implementability. Based on our design specifications, using DNA inversion is the most stringent requisite to reach the largest number of functions. We believe that this is due to the non-destructive nature of DNA inversion compared to DNA excision, and to the fact that DNA-inversion increases the semantic space accessible to a particular architecture. For example, symmetric functions (e.g. XOR), which are highly complex, are easily designed using DNA inversion and nesting pairs of sites. An important flexibility within the number of promoter, gene and terminator also allows to implement more functions.

To navigate within the millions of architectures, we constructed a web-interface allowing users to obtain all architectures implementing a logic function of interest with the possibility of sorting them according to specific design and biological constraints. This web interface will help biological logic designers to obtain minimized integrase-based logic circuits operating in single-cell. Yet, for each function, myriad architectures with highly-divergent designs are possible. Many will not be functional, and not all possible designs can be tested, making the obtention of working logic devices from the database challenging. To circumvent this issue, we reasoned that characterizing and optimizing a single inversion-based device per NP-class would approximate the characterization of all logic devices from this NP-class. This simplification reduces the number of devices to test to 16 for 3-input logic and to 380 for 4-input logic, of which 322 are implementable using only inversion. We provide an example set of the 16 architectures required for 3-input logic implementation.

For a given architecture, specific integrases, integrase site positions and orientations, and gene expression parts will have to be chosen. As highlighted in our previous work ^12^, integrase sites can affect the behavior of logic devices. Previous characterization of individual components coupled with mathematical modeling of part concatenations could be used to limit the number of trial-and-error circles.

A key contribution of our work is the establishment of a formal language for expressing and interpreting genetic construct functionality. The semantic we created, along with our semantic concatenation rules, support unambiguous interpretation of genetic construct functionality. The reduced set of semantics proposed here could be extended to allow interpretation of larger set of genetic constructs, and support numerous synthetic biology applications.

While our minimization strategies allowed us to reduce the generation time to 1h30 for 4-input logic, this systematic generation method is unlikely to be applied for highest number of inputs due to the enormous computation time required (more than one year for 6 inputs). To obtain logic devices responding to a higher number of inputs, future work might focus on extracting from the database systematic design rules generalizable to N-inputs. Additionally, several characterization and optimization cycles could help determine rules for designing architectures responding to an increasing number of inputs. As an alternative approach, the 2-, 3- and 4-input designs proposed here could be concatenated to generate n-input designs.

This work lays the foundation for the systematic design of single-layer, compact recombinase logic devices operating in single-cell. By providing a user-friendly interface to navigate the design space of recombinase-based logic, we empower researchers and engineers to expand the use and design principles of this highly useful class of logic devices.

## Acknowledgments

We thank the synthetic biology group and members of the CBS and LIRMM for fruitful discussions.

## Funding

Support was provided by an ERC Starting Grant “Compucell”, the INSERM Atip-Avenir program and the Bettencourt-Schueller Foundation. S.G. was supported by a Ph.D. fellowship from the French Ministry of Research and the French Foundation for Medical Research (FRM) FDT20170437282. The CBS acknowledges support from the French Infrastructure for Integrated Structural Biology (FRISBI) ANR-10-INSB-05-01.

## Author contributions

S.G. and J.B. designed the project. G.P., F.U. and M.L. designed the RECOMBINATOR algorithm. G.P. wrote the RECOMBINATOR algorithm and implemented the web server. S.G. and G.P. analyzed the data. S.G., G.P. and J.B. wrote the manuscript. All authors approved the manuscript.

## Competing interests

The authors declare no competing interests.

### BOX1: DEFINITIONS

<u>Structure:</u> a sequence of bracket and parenthesis corresponding to recombinase site arrays.

<u>Semantic:</u> gene expression function. The elementary semantics are: promote, terminate, encode, neutral. All sequences have a forward and reverse semantic, written as forward semantic / reverse semantic.

<u>Functional structure:</u> a structure with a semantic between each bracket and parenthesis. A functional structure is composed of 2N brackets and parentheses and 2N+1 semantics, with N the number of inputs.

<u>Part:</u> a promoter, gene or terminator.

<u>Element:</u> composition of parts enabling the “simplest” implementation of a specific forward and reverse semantic. These elements are selected to be composed from a reduced number of parts, to avoid facing promoters, and to reduce the space between a promoter and its transcribed gene.

<u>Architecture:</u> functional structure in which semantics have been replaced by their corresponding elements.

<u>Biological device:</u> an architecture with assignation of input to the parenthesis (integrase sites).

<u>Boolean function:</u> a function with binary variables and a binary output which can be expressed as a propositional formula.

<u>P-class:</u> permutation class, a set of Boolean functions equivalent by permutation of inputs.

<u>NP-class:</u> negation and permutation class, a set of Boolean functions equivalent by permutation and negation of inputs.

<u>Non-redundant:</u> a Boolean function is non-redundant if it cannot be simplified into a Boolean function with a reduced number of inputs. A non-redundant functionalized structure or architecture implement a Boolean function which cannot be simplified into a Boolean function with a number of inputs lower than the number of recombination pairs.

<u>Symmetric:</u> an object is symmetric if it is the reverse complement of itself.

<u>Chiral:</u> two objects (e.g. architectures, structures, functional structures, devices) are chirals if one object is the reverse complement of the other one.

<u>Irreducible:</u> a functionalized structure is irreducible if changing a semantic to a semantic implemented with a reduced number of parts changes the implemented Boolean function (or P-class).

<u>Atomic recombination:</u> an atomic recombination (either excision or inversion) corresponds to a recombination which do not contain other recombination inside it.

## Methods

### 1 Algorithm for architecture generation

The generation algorithm is summarized in Fig S3. To generate the architecture database, we first generated all the structures. Structures correspond to well-balanced sequences of two sets of parentheses ([ ] and ( )) corresponding therefore to all the Dyck word of size 2N (with N the number of inputs) with the two sets of parentheses. For non symmetric structures, we removed one of the two structures of a chiral pair.

The next step of the algorithm consists of assigning a semantic between parentheses. We defined variables as the space between the parentheses, each structure is composed of 2N+1 variables. We defined a semantic domain for each variable, corresponding to a list of semantic which can be assigned to this variable. Domains are composed of either all possible 26 semantics or a reduced set defined to reduce the generation of reducible functionalized structures (see supplementary text 3.2).

For semantic assignation, we used a backtracking algorithm^28^. Briefly, a semantic is assigned to a variable, constraints are evaluated to determine if the semantic is useful. If a constraint is violated, a new track is followed, then the next semantic of the domain is assigned and tested. If no constraint is violated, the assignation algorithm pass to the following variables. All constraints are defined in supplementary text 3.2, the objective of these constraints being that each semantic in both forward and reverse orientations is useful in at least the functionalized structure or one of the derived structures.

When a complete assignation is obtained, the irreducibility of the functionalized structure is tested. To check it, each semantic is replaced with a less complex semantic (see supplementary text 3.2), as an example, fGP is replaced by fG, fP and fN. If the obtained structures implement the same logical function than previously, the initial functionalized structure is reducible.

The truth tables of irreducible functionalized structures are computed by generating all derived structures and computing the semantic of each structure and derived structure. By arbitrarily assigning each parenthesis to an input, one possible Boolean function is obtained and its redundancy can be tested. If the Boolean function cannot be simplified into a function with a reduced number of inputs, the functionalized structure as being non-redundant is saved with its associated Boolean function corresponding to one of the Boolean functions of the implemented P-class.

We implemented the generation algorithm in C++ 17 with Boost library (code available in git directory: https://bitbucket.org/Guigui_PL/genetixV2). We performed the generation for up to four inputs using a high performance computer with a processor Intel Xeon E5-2680 v4, 2.40GHz, 14 physical cores, and with 128Go RAM DDR4 2400Mhz. The generation took around 1h30 for 4 inputs.

### 2 Database creation and Recombinator web-interface

The database construction is decoupled from the generation as the used high performance computer does not have any database management system. From the text files containing the functionalized structures, we created a database satisfying the two normalization form^29^ which permit to not have redundancy in the database. We decomposed the database in five tables: Boolean functions, P-classes, semantics, structures and functionalized structures (Fig S10). Each table contains in addition to its primary key the filtering criteria depending on this key, such as for the structure table, the number of excisions and number of inversions.

We implemented this database generation algorithm in C++ 14 with Boost and libpq++ libraries. The database management system is PostgreSQL (code available in git directory: https://bitbucket.org/Guigui_PL/genetix/src).

The web interface is built with the PHP Framework Yii2. In the web-interface, users fill in a Boolean function of interest either as formulae of propositional logic or as a binary number corresponding to the truth table. A list of architectures is obtained, architectures can be sorted and filtered according to a list of criteria defined in the following section. Users can select a specific architecture and obtain all corresponding information such as the derived architectures.

### 3 Analyse of the architecture database using python scripts

From the database of 19 millions of architectures, a file was generated for each P-class: an example of function, the total number of architectures implementing this P-class and the number of architectures respecting various criteria.

Following is the detailed list of criteria used in this work and their precise definition:

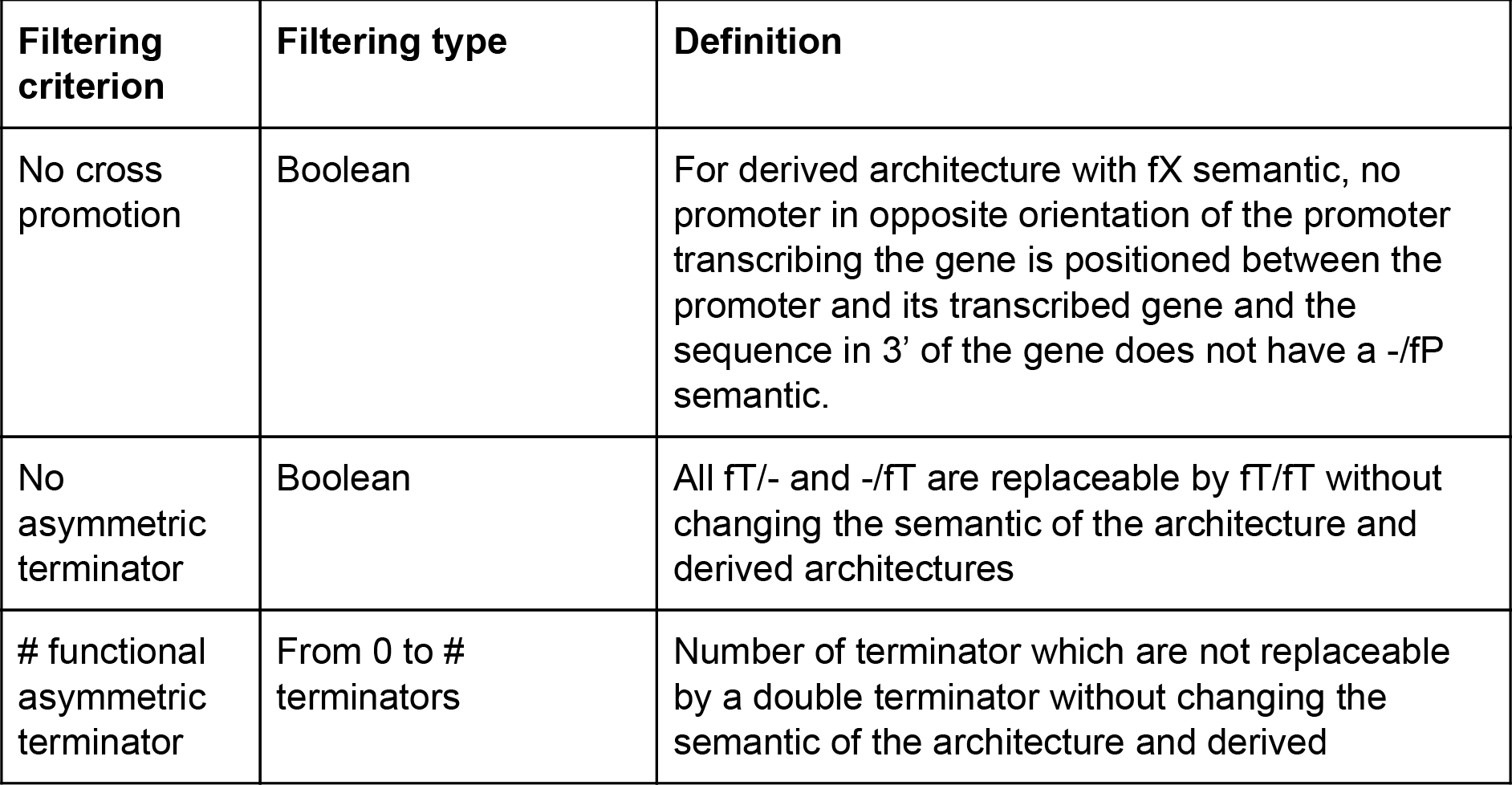
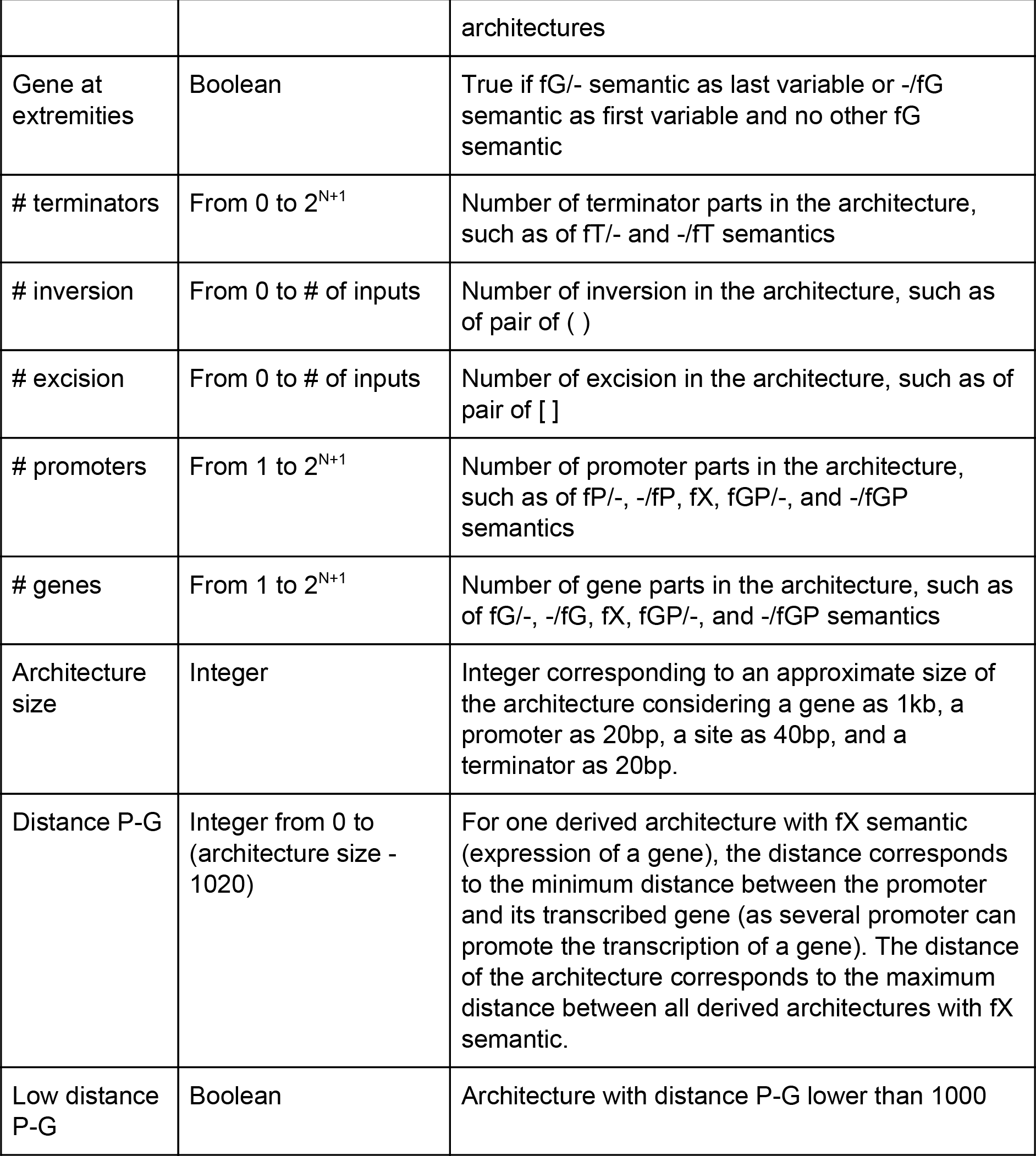

From this file, python scripts were used to extract the percentage table (Fig5) and generated P-class cluster and distribution plots (Fig4, FigS4-S5-S6). For distribution plots, histograms were generated with 20 bins for P-class clusters and 40 bins for histograms corresponding to all P-classes.

Additionally for the database, a file containing the full list of architectures and architecture characteristics was generated. This file was used to generate the two-by-two constraint tables (FigS7-S8-S9) using an automated python script.

### 4 Selection of architectures for the 16 3-input NP-classes

To select one architecture based on inversion per NP-class, the web interface was used. We first generated the list of NP-classes and associated P-classes with one Boolean function per P-classes. From this list, we randomly selected a single Boolean function per NP-class and searched in the database for architectures implementing this Boolean function. We sorted this list of architectures according to the number of excision in decreasing order to select architectures based on inversion only. We then sorted architectures according to the following two boolean constraints: no cross promotion and the low distance between the promoter and the gene. We selected one or several architectures following as much as possible these two criteria and having the lowest number of parts. This selection is not absolute and for most Boolean functions corresponding to one NP-class several architectures existand could be selected.

## Notes

#### Summary of Updates

Correction of Figure 8.

