## Supplementary information for "Exploring the design space of recombinase logic circuits"

### Supplementary Information: Exploring the design space of compacted recombinase logic circuits.

Sarah Guiziou*[†1#], Guillaume Perution-Kihli[†2], Federico Ulliana[2], Michel Leclere[2], and Jerome Bonnet*[1]

[1]Centre de Biochimie Structurale, INSERM U1054, CNRS UMR5048, University of Montpellier, France.
[2]Laboratoire d'Informatique, de Robotique et de Microelectronique de Montpellier (LIRMM). CNRS UMR 5506, University of Montpellier, France.
[†]These authors contributed equally to this work
[#]Current address: Department of Biology, University of Washington, Seattle, Washington 98195, USA

These supplementary materials contain:

-Supplementary Tables S1 to S4.
-Supplementary Figures S1 to S10.
-Supplementary Texts.

### 1 Supplementary Tables

| **Design Specifications** | **Motivations** |
|---|---|
| One pair of sites/integrase | - Reduce problems of non-specific recombination.<br>- Reduces genetic instability.<br>- Reduces difficulties to synthesize.<br>- Reduces the size of the circuit. |
| One integrase by input | - Reduce the number of orthogonal integrase needed.<br>- Reduces metabolic load to the cell.<br>- Reduces the size of the circuit. |
| Regulation of transcription using promoters and terminators | - Simple set of tools.<br>- Two parts for opposite behaviors. |

**Supplementary Table 1: Specification of our logic design.**

| **5' to 3'** | **fN** | **fP** | **fT** | **fG** | **fGP** | **fX** |
|---|---|---|---|---|---|---|
| **fN** | fN | fP | fT | fG | fGP | fX |
| **fP** | fP | fP | fT | fX | fX | fX |
| **fT** | fT | fP | fT | fT | fP | fX |
| **fG** | fG | fGP | fG | fG | fGP | fX |
| **fGP** | fGP | fGP | fG | fX | fX | fX |
| **fX** | fX | fX | fX | fX | fX | fX |

**Supplementary Table 2: Simplification of all possible concatenations of the six semantics two by two**, the row corresponds to the semantic in 5' and the column the semantic in 3'.

| | 1 input | 2 inputs | 3 inputs | 4 inputs | 5 inputs |
|---|---|---|---|---|---|
| # functions with strictly N inputs | 2 | 10 | 218 | 64,594 | $4.3\ 10^9$ |
| # P-classes with strictly N inputs | 2 | 8 | 68 | 3904 | $3.7\ 10^7$ |
| # NP-classes with strictly N inputs | 1 | 5 | 16 | 380 | 1,227,756 |

**Supplementary Table 3: Number of functions, P-classes, NP-classes for a given number of inputs.**

| | > 1 term | > 2 terms | > 3 terms | > 4 terms | > 5 terms | > 6 terms | > 7 terms |
|---|---|---|---|---|---|---|---|
| > 3.5 literals | 25.1% (223) | 26.7% (210) | 36.1% (155) | 38.7% (93) | 60.6% (33) | 53.3% (15) | 0% (2) |
| > 3 literals | 20.7% (1098) | 21.3% (1065) | 25% (908) | 29.6% (514) | 37.2% (188) | 51.8% (56) | 0% (7) |
| > 2.5 literals | 13.3% (2816) | 13.6% (2750) | 16.7% (2241) | 23.7% (1181) | 39.9% (336) | 53.3% (75) | 30% (10) |
| > 2 literals | 10.4% (3599) | 10.66% (3509) | 13% (2875) | 20.3% (1381) | 36.7% (365) | 50% (80) | 30% (10) |
| > 1.5 literals | 9.64% (3879) | 9.9% (3775) | 12.3% (3047) | 19.7% (1418) | 36.2% (370) | 50% (80) | 30% (10) |
| > 1 literal | 9.6% (3894) | 9.87% (3790) | 12.3% (3053) | 19.7% (1418) | 36.2% (370) | 50% (80) | 30% (10) |

**Supplementary Table 4: Percentage of P-classes not implementable for P-classes of specific number of terms and mean number of literal per terms.** Each cell corresponds to the P-classes with a number of term higher than the corresponding column value and a mean number of literal per terms higher than the corresponding line value. And in each cell is filled with the percentage of not implementable P-classes on these specific P-classes and bellow the number of not implementable P-classes.

#### 2 Supplementary Figures

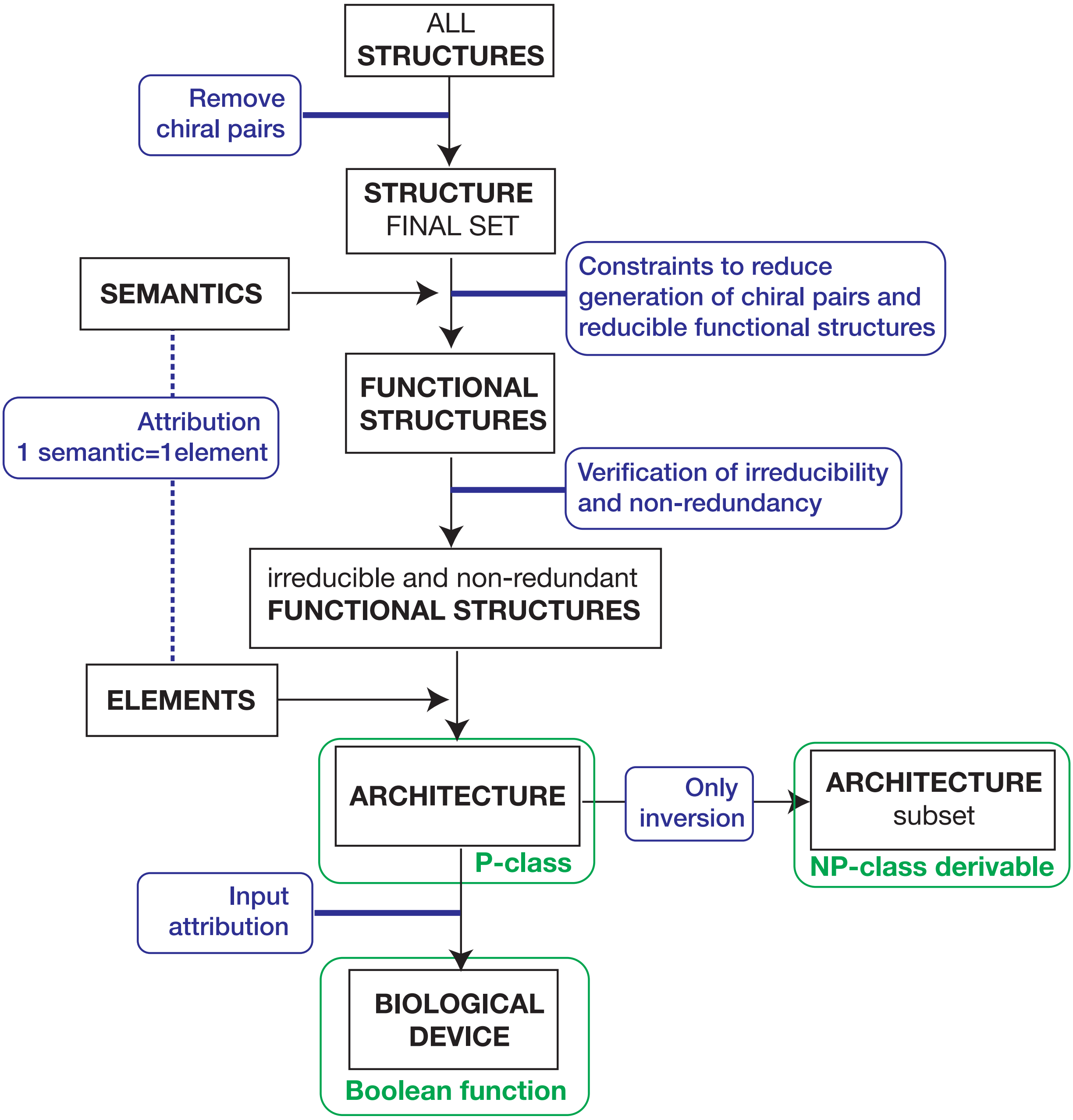


**Supplementary Figure 1**: General workflow of architecture generation.

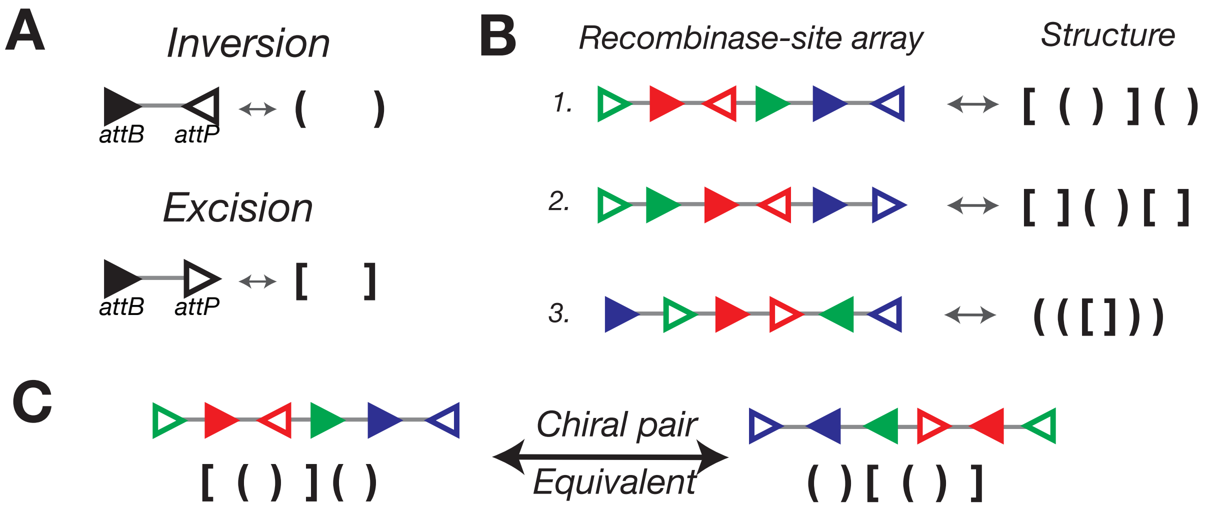


**Supplementary Figure 2**: **Correspondence between recombinase-site array and structure** A - Integrase sites in inversion orientation are represented by parenthesis while integrase sites in excision orientation are represented by brackets. B - Three examples of recombinase-site array and the corresponding structure. In recombinase-site array, each color corresponds to a different input. In the structure, correspondence between parenthesis and input is not specified. C - Example of two chiral structures and their corresponding recombinase-site arrays.

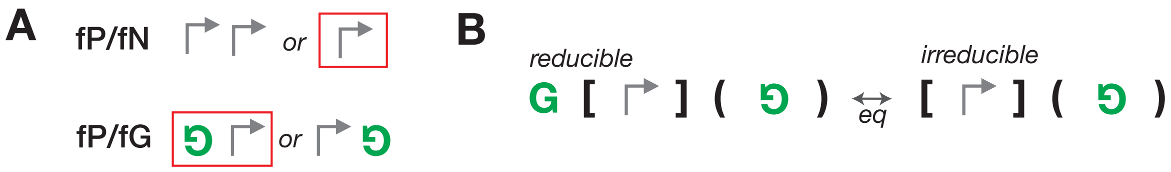


**Supplementary Figure 3**: **Irreducible architecture** A - Example for two semantics of the element selection, two examples of part composition are represented, the one selected as element is surrounded by a red rectangle. For the semantic fP/fN the selection is performed to reduce the number of biological parts while for the semantic fP/fG is performed to avoid cross promotion. B - Two architectures are represented, both implement the same P-class, one being reducible and the other one irreducible.

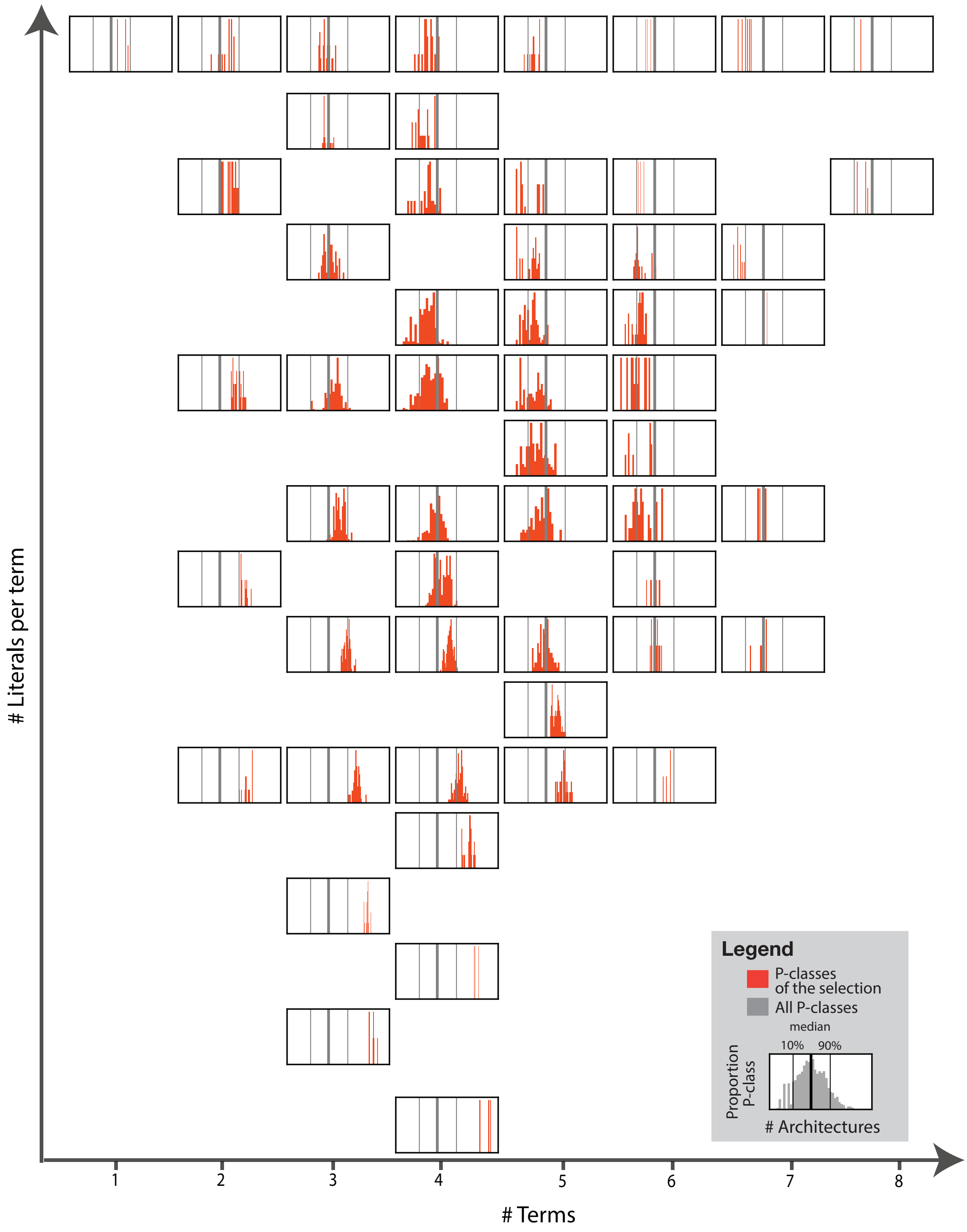


**Supplementary Figure 4**: **Distribution of the number of architecture for the implementation of P-classes with specific number of literal per terms and terms.** Each plot is the distribution of the number of possible architectures (in orange) for a specific set of P-classes; P-classes with a specific number of terms and literals per term (from the lowest in the bottom left to the highest in the top right). These distributions were obtain from a bin of 20. In the legend, the distribution for all P-classes is represented (with a bin of 40). In all graphs, the 10-percentile, the median and the 90-percentile of the distribution of all P-classes is shown in grey.

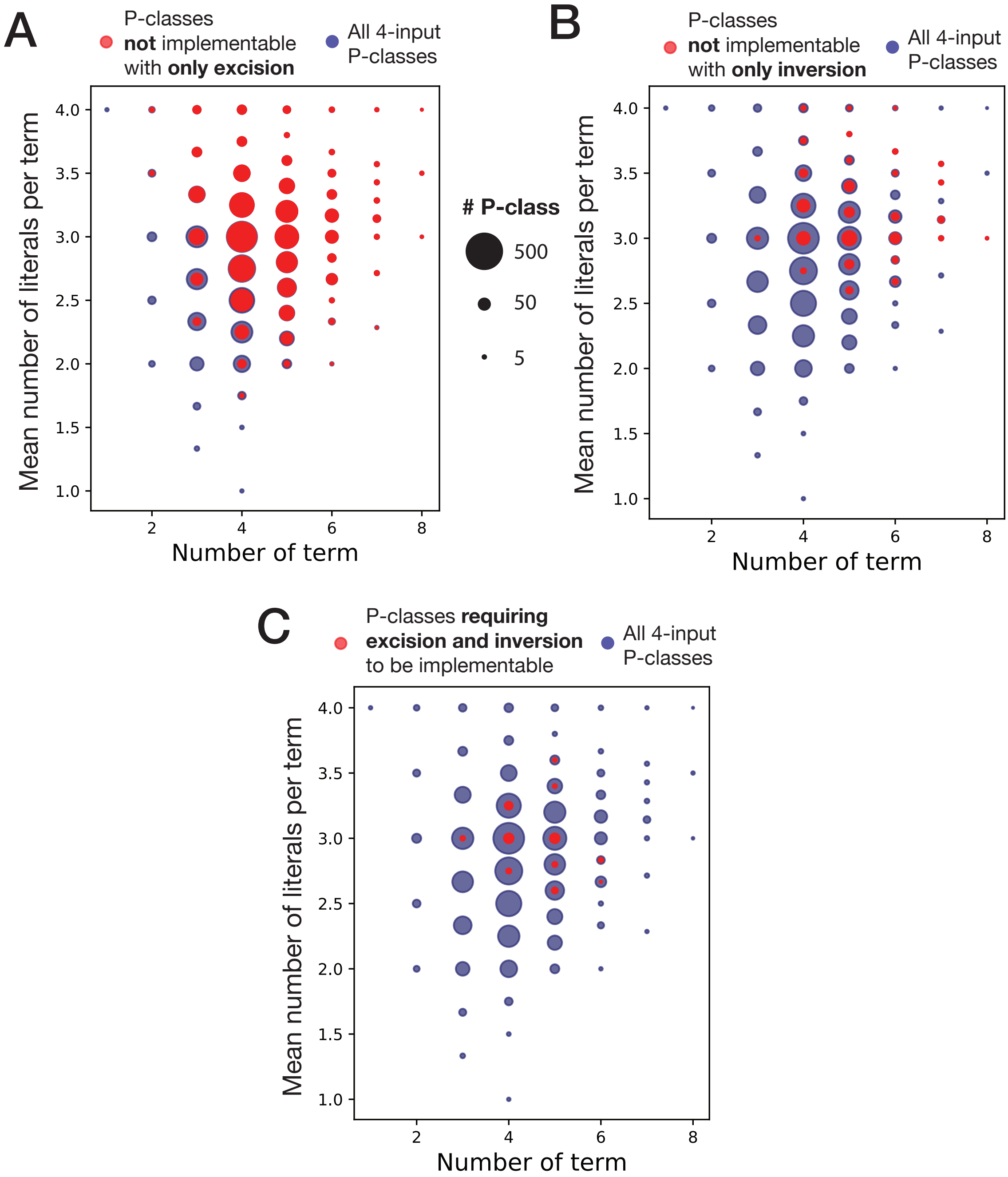


**Supplementary Figure 5**: **Implementability of P-classes with only excision and inversion according to their complexities.** In both plots, P-classes with a specific number of terms and of literals per term are represented in the corresponding position in the plot by a point with the diameter being proportional in logarithm to the number of P-classes. The blue points are for all 4-input P-classes and the red points are for P-classes not implementable with only excision in (A), with only inversion in (B), and P-classes requiring both inversion and excision to be implementable in (C).

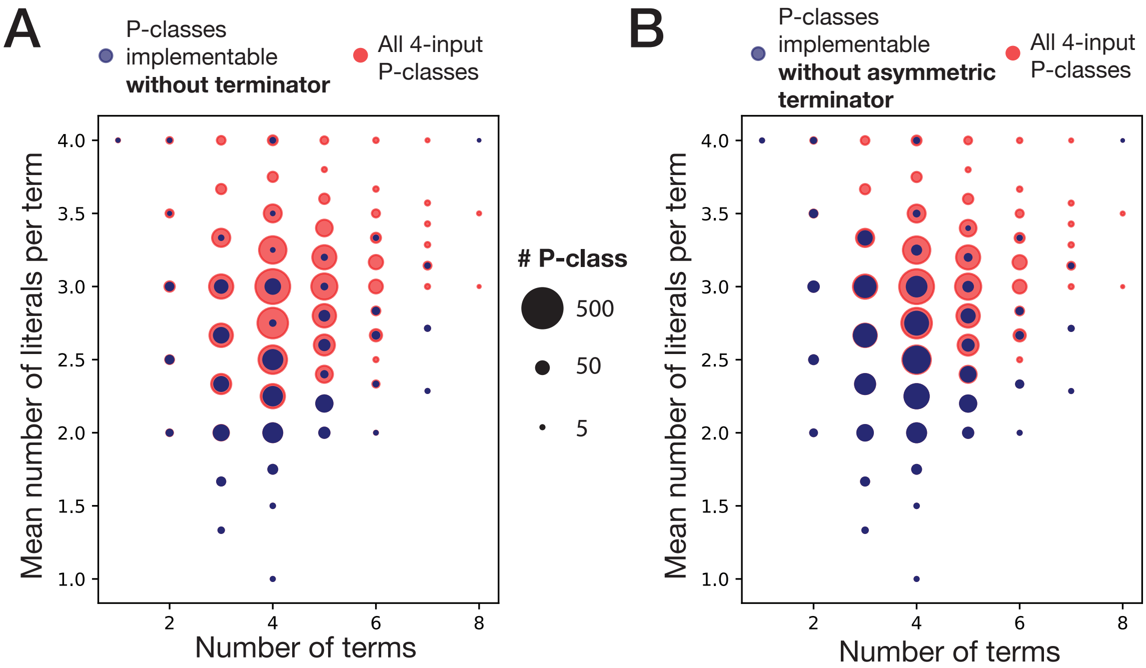


**Supplementary Figure 6**: **Implementability of P-classes without terminator or without asymmetric terminator.** In both plots, P-classes with a specific number of terms and of literals per term are represented in the corresponding position in the plot by a point with the diameter being proportional in logarithm to the number of P-classes. The blue points are for 4-input P-classes implementable without terminators in (A) and without asymmetric terminator in (B), and the orange points are for all 4-input P-classes.

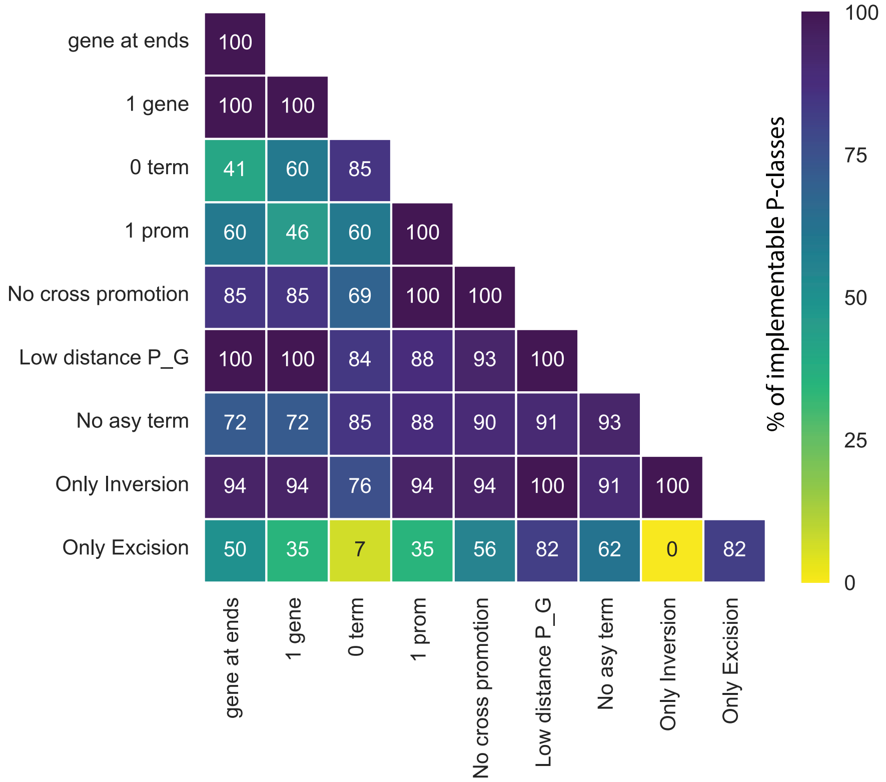


**Supplementary Figure 7**: **Percentage of 3-input P-classes implementable with each constraint and each two-by-two constraint composition.** Each cell corresponds to the percentage of 3-input P-classes implementable with the constraints corresponding to the specific line and column. The color of the cell is related to this percentage, from yellow with 0% to dark blue with 100%. The constraints are explained in the material and methods.

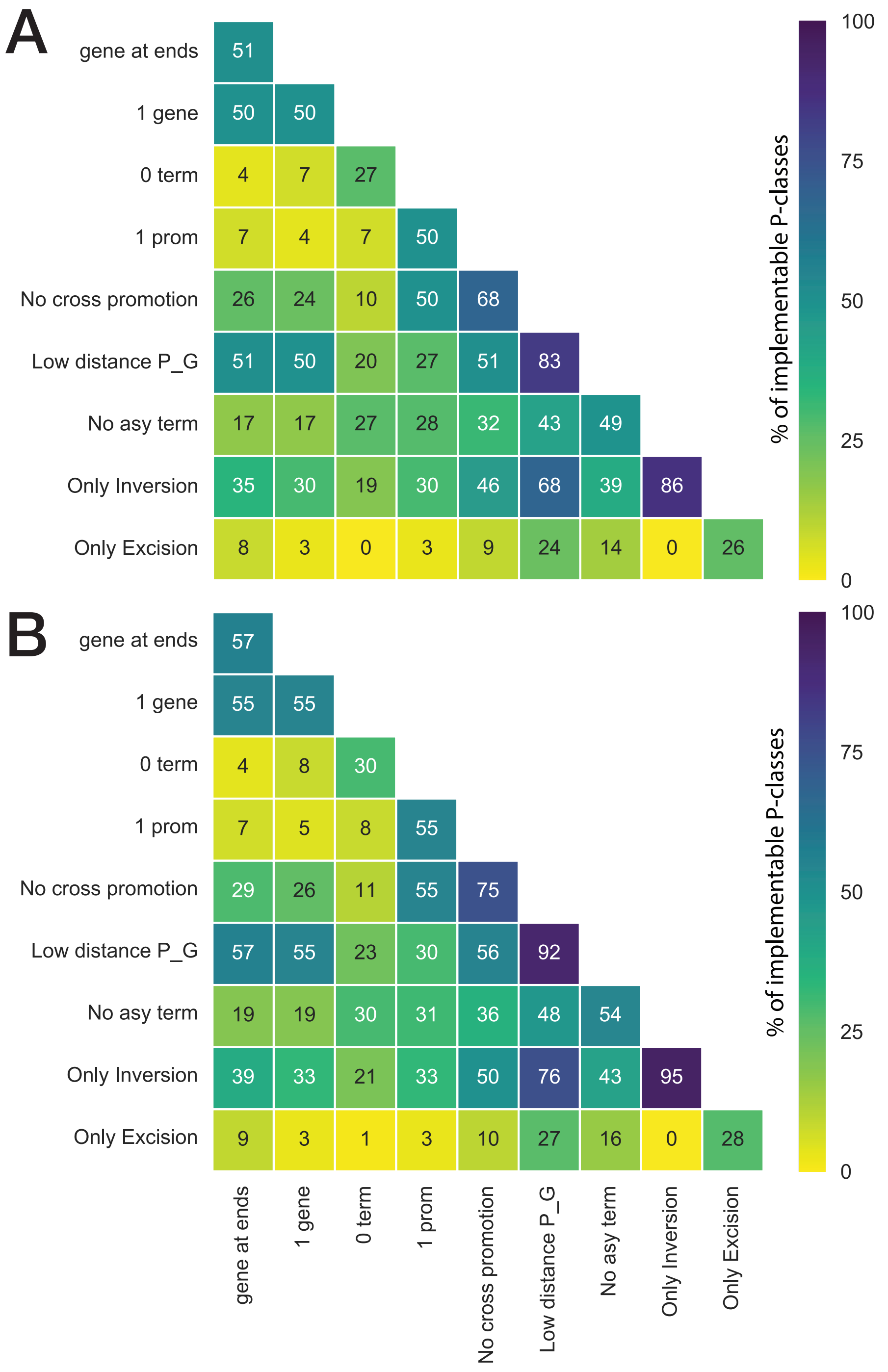


**Supplementary Figure 8**: **Percentage of 4-input P-classes implementable with each constraint and each two-by-two constraint composition.** (A) Percentage on all 4-input P-classes and (B) percentage on the 4-input implementable P-classes. Each cell corresponds to the percentage of implementable P-class with the constraints corresponding to the specific line and column. The color of the cell is related to this percentage, from yellow with 0% to dark blue with 100%. The constraints are explained in the material and methods.

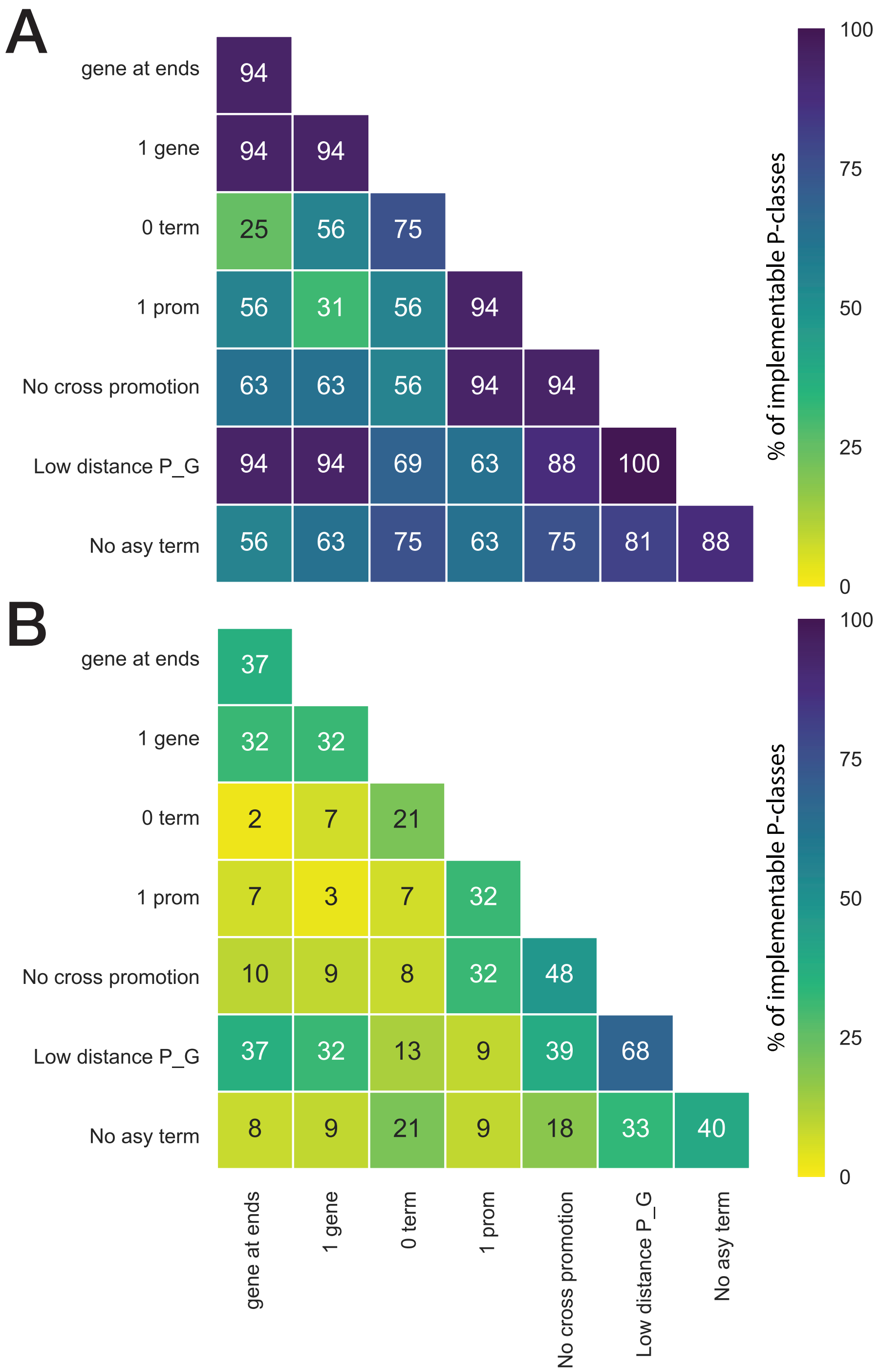


**Supplementary Figure 9**: **Percentage of P-classes implementable with only inversion and with each two-by-two constraint composition.** (A) Percentage on all 3-input P-classes and (B) percentage on all 4-input P-classes. Each cell corresponds to the percentage of implementable P-class with only inversion and the constraints corresponding to the specific line and column. The color of the cell is related to this percentage, from yellow with 0% to dark blue with 100%. The constraints are explained in the material and methods.

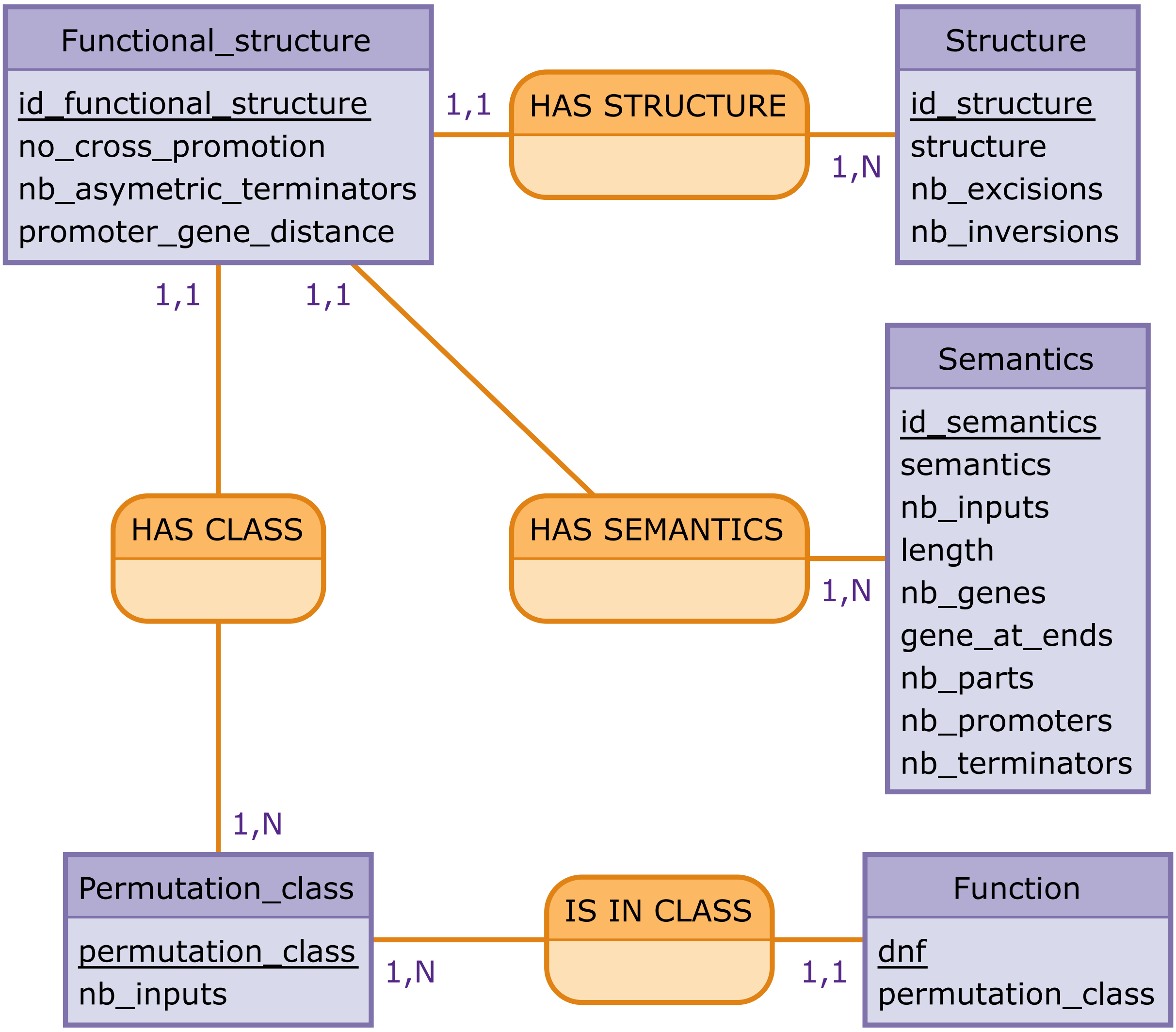


**Supplementary Figure 10**: **Entity-relationship model of the database.** The database is composed of five tables: Boolean functions, P-class, semantics, structure and functionalized structures (corresponding to the purple rectangles). In each rectangle, all variables are listed with the primary key underlined. Their relations are also represented by line with a name specified and either a 1,1 or 1,N relationship.

### 3 Supplementary Texts

#### 3.1 Mathematical formalization of the syntax and semantic of architectures

##### 3.1.1 Syntax of an architecture

An architecture is an expression obtained by combining primitives by concatenation or by nesting an expression $E$ between two marks of an excision or inversion reaction. The *excision* marks are denoted by a pair of square brackets $[E]$ while the *inversion* marks are denoted by a pair of round brackets (). The empty expression is denoted by $\epsilon$ and $EE$ denotes the concatenation. The design language of such architecture is defined by the following grammar:

$$E \ ::= \ \epsilon \ \mid \ S \ \mid \ EE \ \mid \ [\ ] \ \mid \ (\ )$$

$$S \ ::= \ \text{↱} \ \mid \ \text{↲} \ \mid \ \top \ \mid \ \bot \ \mid \ G \ \mid \ \reflectbox{$G$}$$

An architecture without pair of sites is called an element.

##### 3.1.2 Semantics of an architecture

Every biological part has a function, also called semantic. The function of a promoter denoted by fP, is the initiation of transcription. The function of a terminator is to terminate transcription, denoted by fT. The function of a gene is to encode for a RNA/protein, denoted by fG. Also, each part has an orientation which can be either forward or reverse (Fig1C).

The semantic of an architecture (that is, whether it expresses a gene) is a function whose input is a set of active recombinases and the output is a transcriptional value. The transcriptional value is a couple (fF,fR) where fF denotes the semantic of the device in the forward direction and fR denotes the semantic of the device in the reverse direction.

We start by defining the transcriptional semantic of biological parts and elements, as this is independent from any recombinase activation. The transcriptional semantic of a biological part in forward orientation is a couple (f,fN) where f denotes is its semantic (that is, fP or fT or fG) and fN denotes the neutral function meaning that no action is performed. For instance, (fP/fN) is the transcriptional semantic of an architecture reduced to a single forward promoter (Fig1D). In a similar way, the transcriptional semantic of a biological part in reverse orientation is a couple (fN,f) where again f is the function of the biological part.

An element is a concatenation of biological parts. The transcriptional semantic of an elementary sequence is thus given by the left-to-right composition of the semantics of its biological parts in forward orientation together with the right-to-left composition of the semantics of its biological parts in reverse orientation. The composition of semantics of biological parts yields two new semantics, namely the expression of a gene (denoted by fX) and the encapsulation of both encoding and promotion (denoted by fGP). Concretely, the semantic fX is given by the concatenation of promoter with a gene, while the semantic fGP is given by the concatenation of a gene with a promoter.

Importantly, no other function is needed to express the transcriptional semantic of an elementary device. This means that, whatever the size of an element (i.e. the number of its biological parts), its transcriptional semantic is a couple of values (fF,fR) where both fF and fR are one of fN, fP, fT, fG, fGP, fX. Table1C presents the semantics of the composition of the biological parts functions. This means that any element has exactly one among 36 possible transcriptional semantic. Nevertheless, as our objective is too determine the gene expression state of an architecture, all couples (fF,fR) containing at least one fX function are considered as equivalent. In this regard, there are only 26 possible transcriptional semantic for an element.

The transcriptional behaviour of an architecture takes also into account the set of recombinases that are active. Every active pair of sites induces a transformation of the DNA sequence, which can be performed in any order. Once all transformations have been performed, the architecture cannot evolve anymore, and can therefore be seen as an element whose transcriptional semantic can be computed as described before.

The transcriptional semantic of an architecture is a set of couples (A,S) where A is a set of active inputs and S is the semantics of the elementary sequence obtained from the device after all DNA transformations have been performed.

The transcriptional semantic of an architecture can be put in correspondence with Boolean functions. Indeed, a transcriptional semantic can be seen as a truth table where each line corresponds to a couple (A,S). An input that belongs to the set A is active (1) and non active (0) otherwise. Then, the semantic S denotes either gene expression (1) if and only if it contains the behaviour fX.

#### 3.2 Details on the constraints used to reduce the generation of irreducible architectures.

##### 3.2.1 Utility of a sequence according to another one.

A semantic $s_1$ is useful after another semantic $s_2$ if there is no less complex semantic $s_1'$ of $s_1$ such that $s_2.s_1 \equiv s_2.s_1'$. And a semantic $s_1$ is useful before another semantic $s_2$ if there is no less complex semantic $s_1'$ of $s_1$ such that $s_1.s_2 \equiv s_1'.s_2$.

$s_1$ is less complex than $s_2$ if all the corresponding parts of $s_1$ are strictly included in the corresponding parts of $s_2$.

**Example** We defined $s_1 = fP/fN$ and $s_2 = fT/fN$. $s_1$ is useless before $s_2$ because $s_1.s_2 = fT/fN$ and therefore, it exists $s_{1'} = fN/fN$ such that $s_{1'}.s_2 = fT/fN$. But $s_1$ is useful after $s_2$ because there exists no $s_{1'}$ less complex that $s_1$ such that $s_2.s_1 \equiv s_2.s_1'$.

**Constraints** With this concept of utility, we create two constraints, one to check the utility of a semantic after another one, and a second one to check the utility of a semantic before another one, for each variable. A semantic is useful if, on one derived structure, there is an utility in the forward direction and on another one derived structure (not necessarily the same as before), there is an utility in the reverse direction.

All cases of utility are presented in the table S5 of this section. There is only the forward semantics : for the reverse semantics, the table must be read in the other direction.

To know if a semantic is useful before another one, we read the row and after the column. For example, if $s_1 = fP$ and $s_2 = fT$, to check if $s_1$ is useful before $s_2$, we read the row $fP$ and the column $fT$ : in the bottom-left corner of the cell, we can read $fN$, so there exists a $s_{1'} = fN$ such that $s_1.s_2 \equiv s_{1'}.s_2$. If the cell was blank, $s_1$ would be useful before $s_2$.

To know if a semantic is useful after another one, we read the row before the column. For example, if $s_1 = fGP$ and $s_2 = fP$, to check if $s_1$ is useful before $s_2$, we read the column $fGP$ and the row $fP$ : in the top-right corner of the cell, we can read $fG$, so there exists a $s_{1'} = fG$ such that $s_1.s_2 \equiv s_{1'}.s_2$. If the cell was blank, $s_1$ would be useful before $s_2$.

| $s_1.s_2$ | fN | fP | fT | fG | fGP | fX |
|---|---|---|---|---|---|---|
| fN | | | | | | |
| fP | | fN \ fN | fN \ | | \ fG | fN \ fG |
| fT | | fN \ | fN \ fN | \ fN | \ fP | fN \ |
| fG | | | \ fN | fN \ fN | fN \ fP | fN \ |
| fGP | | fG \ fN | fG \ | fP \ | fP \ fG | fN \ fG |
| fX | | \ fN | \ fN | fP \ fN | fP \ fN | fN \ fN |

**Supplementary Table 5: utilities of a semantic concatenated to another one**

##### 3.2.2 Utility of prefix and suffix semantics.

The principle here is similar to the one described before. Instead of compare a semantic to a semantic before or after another one, we compare it to the prefix semantic or the suffix semantic of the variable in the derived functional structures.

The prefix semantic of a variable is the concatenation of all semantics before this variable. And the suffix semantic is the concatenation of all semantics after this variable.

The table S5 of this section must be read in the same way as the table S6.

| $Sem_{column}.Sem_{row}$ | fN | fP | fT | fG | fGP | fX |
|---|---|---|---|---|---|---|
| fN | | | \ fN | \ fN | \ fP | |
| fP | fN \ | fN \ fN | fN \ | | \ fG | fN \ fG |
| fT | fN \ | fN \ | fN \ fN | \ fN | \ fP | fN \ |
| fG | | | \ fN | fN \ fN | fN \ fP | fN \ |
| fGP | fG \ | fG \ fN | fG \ | fP \ | fP \ fG | fN \ fG |
| fX | | \ fN | \ fN | fP \ fN | fP \ fN | fN \ fN |

**Supplementary Table 6: utilities prefix and suffix.**

##### 3.2.3 Domain definitions

By default, each variable has in its domain all the 26 semantics. But, in some cases, we can reduce the domain of some variables.

**Atomic inversion and excision** A pair of sites is atomic if there is no site between them.
In an atomic excision, we do not assign the $fN/fN$ semantic because. Indeed, after excision, the semantic will remained identical ($fN/fN$) leading to the implementation of a redundant Boolean function.
Identically for atomic inversion to avoid redundant Boolean function, we do not assign symmetric semantics as they are not modified by inversion (fN/fN, fP/fP, fG/fG, fT/fT, fGP/fGP, fX).

**Expressed semantic** The semantic fX can only be put in an excision. Otherwise, the gene would be always expressed on all derived structures leading to the implementation of the True Boolean function which is redundant.

**Semantics at the ends of a functional structures** As a consequence of the prefix and suffix constraints, we only assign four semantics in the first variable (fN/fN, fP/fN, fN/fG, fP/fG) and four in the last variable (fN/fN, fN/fP, fG/fN, fG/fP). Indeed, the prefix of the first variable is always fN/fN to respect the previously defined constraints these are the only semantics that have a prefix utility after fN/fN. The same applies with the suffix of the last variable.

#### 3.3 Rules permitting to simplify a concatenation of semantics, permitting therefore to determine the gene expression state.

Transcriptional parts are concatenated to form transcriptional sequences. We defined a set of rules to determine the semantics of sequences. As any forward and reverse semantics can be considered separately, the following properties are defined considering a single orientation of the construct. For simplification, the properties are written in the forward orientation, from 5' to 3'.

The semantic of a transcriptional sequence corresponds to the concatenation of the semantics of each part of the sequence. Indeed, to determine the semantics of a concatenation of transcriptional parts, we use a step-wise iterative process in which semantics are composed two by two.

This concatenation can be simplified with the following rules in a reduced set of six elementary semantics. These six semantics correspond to the four semantics described previously (fP, fT, fG, and fN) plus the semantics corresponding to the expression of a gene: fX and the composition of fG followed by fP: fGP. As the two-by-two concatenation of the four basic semantics leads to one of these six semantics, this set of semantics is complete (detailed below).

Rules:

**(1) Non-commutativity**: Concatenation of semantics is not commutative as parts concatenated in a different order do not lead to the same semantic. As an example, PF-GF permits expression of the gene, therefore encoding the semantic fX, which is not the case for GF-PF.

**(2) Neutrality**: A sequence without any activity in a particular orientation does not affect other sequences placed in the same orientation (i.e. fN is neutral to other semantics similarly oriented).
**(3) Assimilation of fX**: The semantic of gene expression, fX, assimilates all other semantics. In others words, the composition of the fX semantic with another semantic is simplifiable to fX. In this work, we aim at defining if a construct leads to expression of a gene or not and the composition of fX with another semantic does not affect the fX semantic.
**(4) Idempotent**: all semantics are idempotent (an operation has the same effect even if applied multiple times), as we consider that the concatenation of two similar parts is equivalent to a single part.

Others rules are due to the mechanism of gene expression. A gene is expressed if it is transcribed by RNA polymerase; consequently, a promoter needs to be positioned upstream without a terminator positioned between the promoter and the gene. This mechanism can be assimilated to a flow that is opened by the promoter, stopped by the terminator, and the system is ON when the flow is at a specific location: the gene.
(5) Expression occurs only if a promoter is placed upstream of a gene without a terminator in between. Such as, the concatenation of the semantic promotion with semantic gene is simplifiable to fX. i.e. fP-fG=fX
(6) A gene followed by a promoter leads to the semantic fGP, as the promoter can be active for a downstream gene and the gene can be expressed by an upstream promoter. Consequently, **fG-fP=fGP**. In this case, we have associativity of the semantics fG and fP.
(7) If a promoter is followed by a terminator, the RNA polymerase flux is blocked by the terminator, consequently, *fP-fT=fT*.
(8) If a terminator is followed by a promoter, the terminator will have no effect on the semantic of the sequence as the terminated transcription will be re-initiated by the promoter, consequently: **fT-fP=fP**.
(9) If a terminator is followed by a gene, the gene cannot be expressed as the RNA polymerase flux is blocked by the terminator; consequently, **fT-fG=fT**.
(10) If a gene is followed by a terminator, the terminator will have no effect on the semantic of the word; indeed, if the previous semantic is fP, it will result in the expression of the gene, consequently: **fG-fT=fG**.

**Mathematical definition of the rules for semantic composition.** For two semantics fA and fB, fA-fB is the concatenation of fA with fB, fA being in 5' and fB in 3'.
*(1) Not commutative: fA-fB≠fB-fA*
*(2) Neutrality of fN: fN-fA=fA and fA-fN=fA*
*(3) Assimilation by fX: fX-fA=fX and fA-fX=fX*
*(4) Isomorphism: fA-fA=fA*
*(5) Condition of gene expression: fP-fA-fG=fX only if fA≠fT or fA=fP*
*(6) Composition of Gene-Promoter: fG-fP=fGP*
*(7) A terminator cancels promotion: fP-fT=fT*
*(8) A promoter cancels termination: fT-fP=fP*

*(9) A terminator block transcription of a following gene: fT-fG=fT*
*(10) A terminator cannot block transcription of a previous gene: fG-fT=fG*

We concatenated the six previously defined semantics two by two. Using the previously defined rules, all concatenations of these six semantics are simplifiable to one of the six semantics (TableS2). Therefore, the set of semantics is complete and our rules are scalable to the concatenation of N transcriptional parts.

#### 3.4 Symmetric functions are implementable with recombinases.

##### 3.4.1 Definition of a symmetric function

**A symmetric function of n variable is a function which is equivalent by any permutation of the n variable.**
As examples:
(1) f(a,b,c)=a.b.c+!a.!b.!c (with !x being the negation of x) is a symmetric function which is equal to 1 if the number of one variable is zero or three. Therefore, it can be written as $S^3_{0,3}$(a,b,c), a,b and c being the variable of the symmetric function.
(2) f(a,b,c)=a.!b.c+!a.b.!c is also a symmetric function. Here the variables are a, !b, c and the function can be written as $S^3_{0,3}$(a,!b,c).

We note symmetric functions as: $S^n_{\{a_k\}}(x_1^{j_1}, x_2^{j_2}, ..., x_n^{j_m})$, where the $j_i$ may take on only the values 0 and 1, and:

$$x_i^{j_i} = x_i \text{ if } j_i = 0 \text{ and } x_i^{j_i} = \neg x_i = !x_i \text{ if } j_i = 1$$

##### 3.4.2 Formal implementation of symmetric functions with recombinases.

Symmetric functions are usually functions with a large number of terms and of literal per terms, so requiring large circuits for implementation if using second-order circuits based on sum-of-products or product-of-sums function expression. However, in electronic, in the 1960 century, methods have been developed to easily and economically realize completely symmetric functions using contact-type gating elements. As well in biology, symmetric functions are easily implementable using recombinases using nested inversion-based elements.

As example, the 2-input XOR gate were realized by Bonnet and colleagues using a single terminator nested inversion-based elements as represented in Figure bellow. In the presence of none of the input, the terminator is blocking the expression of the output gene. In presence of a single input, the terminator is inverted leading to expression of the output gene. In presence of the two input, the terminator is inverted back in its blocking orientation, the output gene is then not express. This design can be generalize for an n number of input for even and similarly odd symmetric functions (Figure S11).

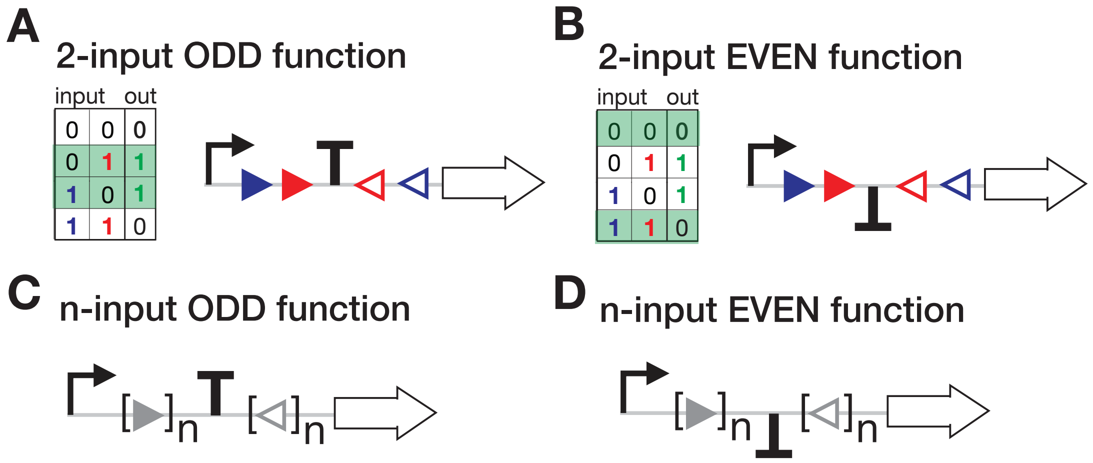


**Supplementary Figure 11**: **Implementation of odd and even symmetric functions using recombinases.** (A) For the implementation of 2-input odd function also called XOR function, Bonnet and colleagues used a terminator nested inversion-based element. In more detail, the device is composed on an asymmetric terminator in blocking orientation surrounded by two nested pairs of integrase sites in inversion orientation. (B) Similarly, for the implementation of the 2-input even function also called NXOR function, Bonnet and colleagues used a terminator nested inversion-based element, however here the asymmetric terminator is in passing orientation in the off state. (C) and (D) These 2-input designs can be generalized to N-input designs by nested N-integrase site pairs in inversion orientation.

##### 3.4.3 All fully implementable clusters of complex functions are composed of symmetric or partially symmetric Boolean functions.

**List of all logic functions with 8 terms and 4 literals per terms:**
ODD PARITY FUNCTION: $S^4_{1,3}$(a,b,c,d)=!a.!b.!c.d + !a.!b.c.!d + !a.b.!c.!d + a.!b.!c.!d + !a.b.c.d + a.!b.c.d + a.b.!c.d + a.b.c.!d
EVEN PARITY FUNCTION: $S^4_{0,2,4}$(a,b,c,d)=!a.!b.!c.!d + !a.!b.c.d + !a.b.!c.d + !a.b.c.!d + a.!b.!c.d + a.!b.c.!d + a.b.!c.!d + a.b.c.d

**List of all logic functions with 7 terms and 4 literals per terms:**
$S^4_{2,4}$(a,b,c,d)=!a.!b.c.d + !a.b.!c.d + !a.b.c.!d + a.!b.!c.d + a.!b.c.!d + a.b.!c.!d + a.b.c.d
$S^3_{1,3}$(a,b,c).!d+$S^3_{2}$(a,b,c).d=!a.!b.c.!d + !a.b.!c.!d + a.!b.!c.!d + !a.b.c.d + a.!b.c.d + a.b.!c.d + a.b.c.!d
$S^4_{1}$(a,b,c,d)+$S^3_{2}$(b,c,d).a==!a.!b.!c.d + !a.!b.c.!d + !a.b.!c.!d + a.!b.!c.!d + a.!b.c.d + a.b.!c.d + a.b.c.!d
$S^4_{0,4}$(a,b,c,d)+$S^2_{1}$(a,b).$S^2_{1}$(c,d)=!a.!b.!c.!d + !a.b.!c.d + !a.b.c.!d + a.!b.!c.d + a.!b.c.!d + a.b.!c.!d + a.b.c.d
$S^4_{0,2}$(a,b,c,d)=!a.!b.!c.!d + !a.!b.c.d + !a.b.!c.d + !a.b.c.!d + a.!b.!c.d + a.!b.c.!d + a.b.!c.!d

**List of all logic functions with 8 terms and 3.5 literals per terms:**

$S^4_{1,3,4}$(a,b,c,d)=!a.!b.!c.d + !a.!b.c.!d + !a.b.!c.!d + a.!b.!c.!d + b.c.d + a.c.d + a.b.d + a.b.c

$S^2_1$(c,d).!a.!b+c.d.$S^2_1$(a,b)+$S^2_{0,1}$(c,d).a.b+!c.!d.$S^2_{1,2}$(a,b)=!a.!b.!c.d + !a.!b.c.!d + !a.b.c.d + a.!b.c.d + b.!c.!d + a.!c.!d + a.b.!c + a.b.!d

$S^3_{0,2,3}$(a,b,c).!d+$S^3_{1,3}$(a,b,c).d=!a.!b.!c.!d + !a.!b.c.d + !a.b.!c.d + a.!b.!c.d + b.c.!d + a.c.!d + a.b.!d + a.b.c

$S^3_{0,2}$(b,c,d).!a+$S^3_{0,1,3}$(b,c,d).a =!a.!b.c.d + !a.b.!c.d + !a.b.c.!d + a.b.c.d + !b.!c.!d + a.!b.!c + a.!b.!d + a.!c.!d

$S^4_{0,1,3}$(a,b,c,d)=!a.b.c.d + a.!b.c.d + a.b.!c.d + a.b.c.!d + !a.!b.!c + !a.!b.!d + !a.!c.!d + !b.!c.!d

#### 3.5 List of 4-input P-classes with a single possible architectures.

Six 4-input P-classes are implementable with a single possible architectures, following an example of logic function in a Quine McKluskey form for each of these 6 P-classes.

f1=!a.b.c.d + a.!b.!c + a.!b.!d + a.!c.!d

f2=a.b.c.d + !a.!b.!d + !a.!c.!d + !b.!c.!d

f3=a.!b.!c + a.!b.!d + a.!c.!d + b.c.d

f4=!a.!b.!d + !a.!c.!d + !b.!c.!d + a.b.c

f5=a.!b.!c + a.!b.!d + a.!c.!d + !a.c.d + !a.b.d + !a.b.c

f6=!a.!b.!d + !a.!c.!d + !b.!c.!d + b.c.d + a.c.d + a.b.d

#### 3.6 4-input P-class with the maximum number of architectures.

Following an example of logic function of the P-class with the maximum number of architectures (1.4 millions).

f=a+b+!c+!d
